# A visual tour of carbon export by sinking particles

**DOI:** 10.1101/2021.02.16.431317

**Authors:** Colleen A. Durkin, Ken O. Buesseler, Ivona Cetinić, Margaret L. Estapa, Roger P. Kelly, Melissa Omand

**Affiliations:** Moss Landing Marine Laboratories, Moss Landing, CA, USA; Woods Hole Oceanographic Institution, Woods Hole, MA, USA; Universities Space Research Association and the Ocean Ecology Laboratory at NASA/Goddard Space Flight Center, Greenbelt, Maryland, USA; University of Maine, Orono, ME, USA; Skidmore College, Saratoga Springs, NY, USA; University of Rhode Island, Graduate School of Oceanography, Narragansett, RI, USA

**Keywords:** biological pump, carbon export, sediment traps, fecal pellet, aggregate, salp, particles

## Abstract

To better quantify the ocean’s biological carbon pump, we resolved the diversity of sinking particles that transport carbon into the ocean’s interior, their contribution to carbon export, and their attenuation with depth. Sinking particles collected in sediment trap gel layers from 4 distinct ocean ecosystems were imaged, measured, and classified. The size and identity of particles was used to model their contribution to particulate organic carbon (POC) flux. Measured POC fluxes were reasonably predicted by particle images. Nine particle types were identified, and most of the compositional variability was driven by the relative contribution of aggregates, long cylindrical fecal pellets, and salp fecal pellets. While particle composition varied across locations and seasons, the entire range of compositions was measured at a single well-observed location in the subarctic North Pacific over 1 month, across 500 m of depth. The magnitude of POC flux was not consistently associated with a dominant particle class, but particle classes did influence flux attenuation. Long fecal pellets attenuated most rapidly with depth whereas certain other classes attenuated little or not at all with depth. Small particles (<100 *μ*m) consistently contributed ∼5% to total POC flux in samples with higher magnitude fluxes. The relative importance of these small particle classes (spherical mini pellets, short oval fecal pellets, and dense detritus) increased in low flux environments (up to 46% of total POC flux). Imaging approaches that resolve large variations in particle composition across ocean basins, depth, and time will help to better parameterize biological carbon pump models.

## Introduction

Particles that sink out of the surface ocean are an important component of the global carbon cycle because they transport carbon away from the atmosphere and into the deep ocean (Martin 1990; Maier-Reimer et al. 1996). Sinking particles are one component of the biological carbon pump, together with physical subduction of carbon to depth and excretion/egestion of carbon by vertically migrating zooplankton. The amount of carbon transported by the biological carbon pump is highly variable and difficult to accurately constrain (5-12 Pg-C year^-1^, Boyd and Trull 2007; Henson et al. 2011), leading to uncertainties in carbon cycle models. These uncertainties are caused, in part, by the variety of ecological mechanisms that generate particles and the variety of pathways that carbon can take on its way to the deep ocean. After phytoplankton fix carbon in the surface, these cells can sink directly or they can be consumed by zooplankton and egested in fecal pellets, which have highly variable carbon densities and sinking speeds (Alldredge et al. 1987; Urban-Rich et al. 1998; Turner 2015). Alternatively, phytoplankton carbon can be incorporated into aggregates and other amorphous detritus (Alldredge and Gotschalk 1989; Passow et al. 1994). To further complicate the process, carbon can transition from one particle pool to another as it sinks through the mesopelagic and interacts with grazers and microbes. Each of these particle types are thought to transport carbon with differing efficiencies through the mesopelagic (Buesseler et al. 2007). To better constrain the biological carbon pump, mechanistic models must predict the flow of carbon through these pathways and particle types with depth (Siegel et al. 2014). As we understand more about the types of particles sinking through the water column, the accuracy of these models will improve, and we will be able to better predict the magnitude of carbon flux and its attenuation with depth.

The perceived importance of different particle types in exporting carbon has changed over approximately 50 years of study. For example, Turner (2002) reviewed early studies of the biological carbon pump that attributed most carbon export to zooplankton fecal pellets. As more studies were conducted, fecal pellets were found to attenuate rapidly with depth and the dominant flux contributor was instead attributed to phytodetrital aggregates. After additional years of study, the perceptions have shifted again (Turner 2015). Now it is understood that no single paradigm describes the biological pump across the global ocean, except perhaps that particle composition is highly variable in time and space, as are the effects of those particle types on POC export and attenuation (Passow and Carlson 2012; Turner 2015; Boyd et al. 2019). Earlier studies of the biological pump focused on the influential role of large sinking particles (Fowler and Knauer 1986), and recent studies have identified an important role for small (<100 *μ*m) and slowly settling (<20 m d^-1^) particles (Alonso-González et al. 2010; Baker et al. 2017; Bol et al. 2018). In spite of their small size, these particles have been observed in mesopelagic sediment traps (Durkin et al. 2015) and the increasing recognition of “particle injection pumps” suggests that they can be efficiently transported out of the surface mixed layer through physical processes (Barth et al. 2002; Omand et al. 2015; Stukel et al. 2017) or produced at depth from disaggregation of larger particles (Briggs et al. 2020). The perceived importance of various processes and particle types have also changed as technology has advanced. For example, episodic fluxes can now be detected by autonomous instruments (Briggs et al. 2011; Estapa et al. 2013; Huffard et al. 2020) but are not captured in steady state models or by many traditional observational designs. These often cryptic events can sometimes account for a large percentage of the total carbon exported in an ecosystem (Smith et al. 2018).

In situ imaging promises to expand observations of sinking particles, and potentially lead to improved estimates of the ocean’s biological carbon pump (Picheral et al. 2010; Bishop et al. 2016; McGill et al. 2016). These approaches assume that the quantity of carbon export can be predicted by the quantity and character of sinking particles observed in sensor imagery. Here, we demonstrate how the biological characteristics of particles collected in sediment traps (identity, size, abundance) can be used to predict the magnitude and attenuation of carbon flux with depth. This study also explores how the composition of sinking particles affects carbon flux across locations, time, and depth. Our ultimate goal is to use imaging tools to provide the quantitative, particle-resolving observations of carbon flux that are needed to generate mechanistic, ecologically-realistic models of the biological carbon pump.

## Materials

### Cruises and sampling platforms

Samples from cylindrical sediment traps with polyacrylamide gel layers were collected at the New England shelf break aboard the R/V Endeavor on 3-7 November 2015 (EN572) and 13-18 June 2016 (EN581). Samples were also collected at two locations in the subtropical North Pacific and one location in the California Current on a transit between Honolulu, Hawaii and Portland, Oregon aboard the R/V Falkor between 24 January-20 February, 2017 (FK170124). Additional samples were collected during the NASA EXPORTS field campaign near Ocean Station PAPA in the subarctic North Pacific during three successive collection periods (“epochs”) between 10 August-12 September 2018 (Supplemental Fig. 1, Table 1; Estapa et al. in review; Siegel in review). Sediment trap tubes were deployed on various platform designs, including neutrally-buoyant sediment traps (Estapa et al. 2020), surface tethered sediment traps (Knauer et al. 1979) and on the base of a Wirewalker profiler (Rainville and Pinkel 2001). The neutrally-buoyant traps carried 4 collection tubes with a diameter of 12.7 cm and height of 70 cm (Estapa et al. 2020). At the New England shelf break, the surface-tethered trap was comprised of gimbaled trap frames (KC Denmark) clipped on a surface-tethered, free drifting array with five collection depths. Each of the 5 trap frames carried four 7 cm diameter 45 cm high collection tubes. In the subarctic Pacific, the surface-tethered trap frames carried four 12.7 cm diameter collection tubes at 5 depths. The Wirewalker trap consisted of one 4-tube trap frame (KC Denmark) tethered by a bungee below the profiling part of the Wirewalker drifting array. Sample depths and durations are indicated in Table 1.

**Figure 1.**
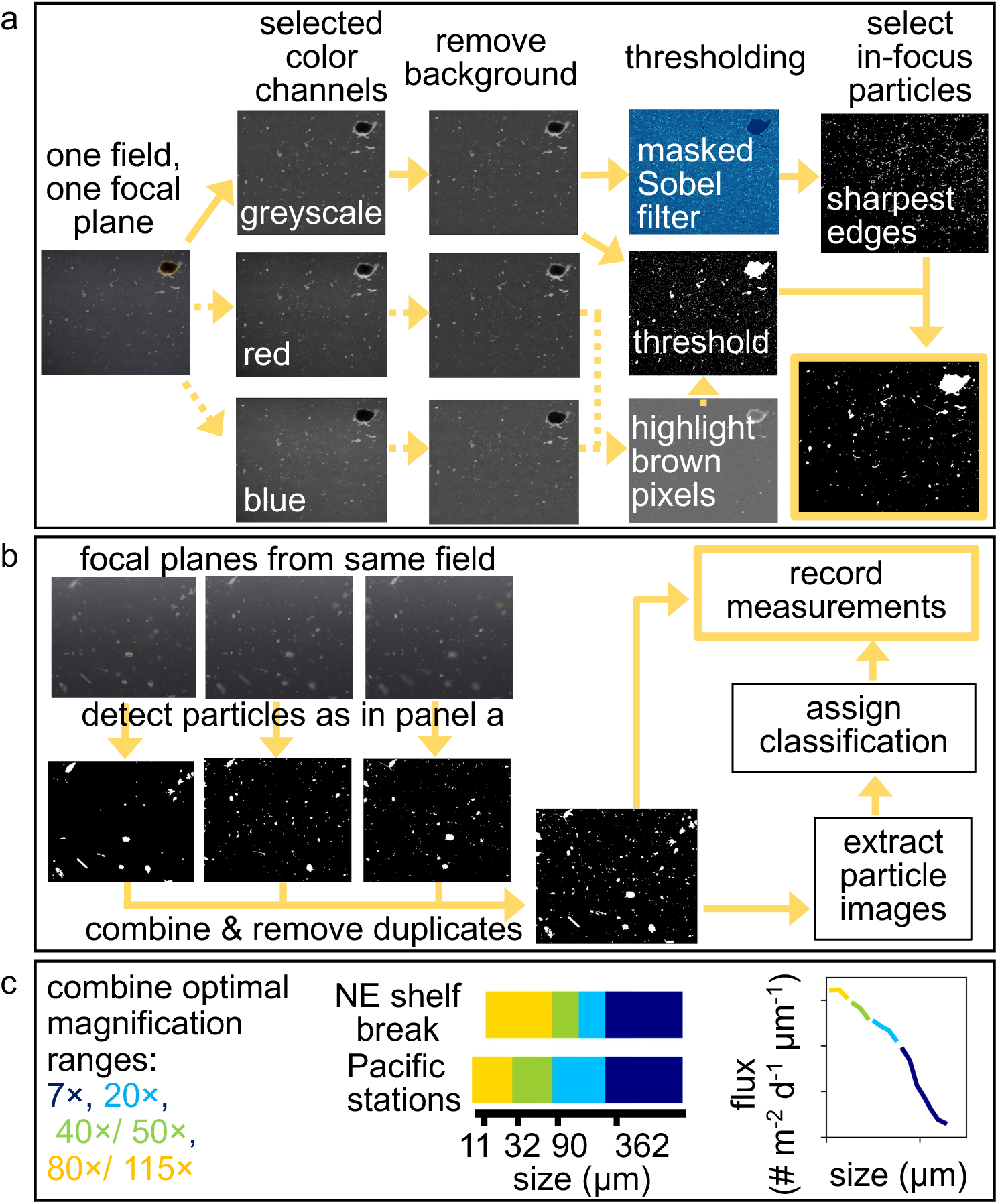
Schematic illustration of image processing methods including a) detection of particles, b) combination of multiple focal planes from the same field of view and collection of particle measurements, and c) combination of flux measurements from optimal magnification ranges. Processing steps for brightfield images are indicated by solid arrows while processing of oblique images also included steps indicated by arrows with dashed lines.

**Table 1.**
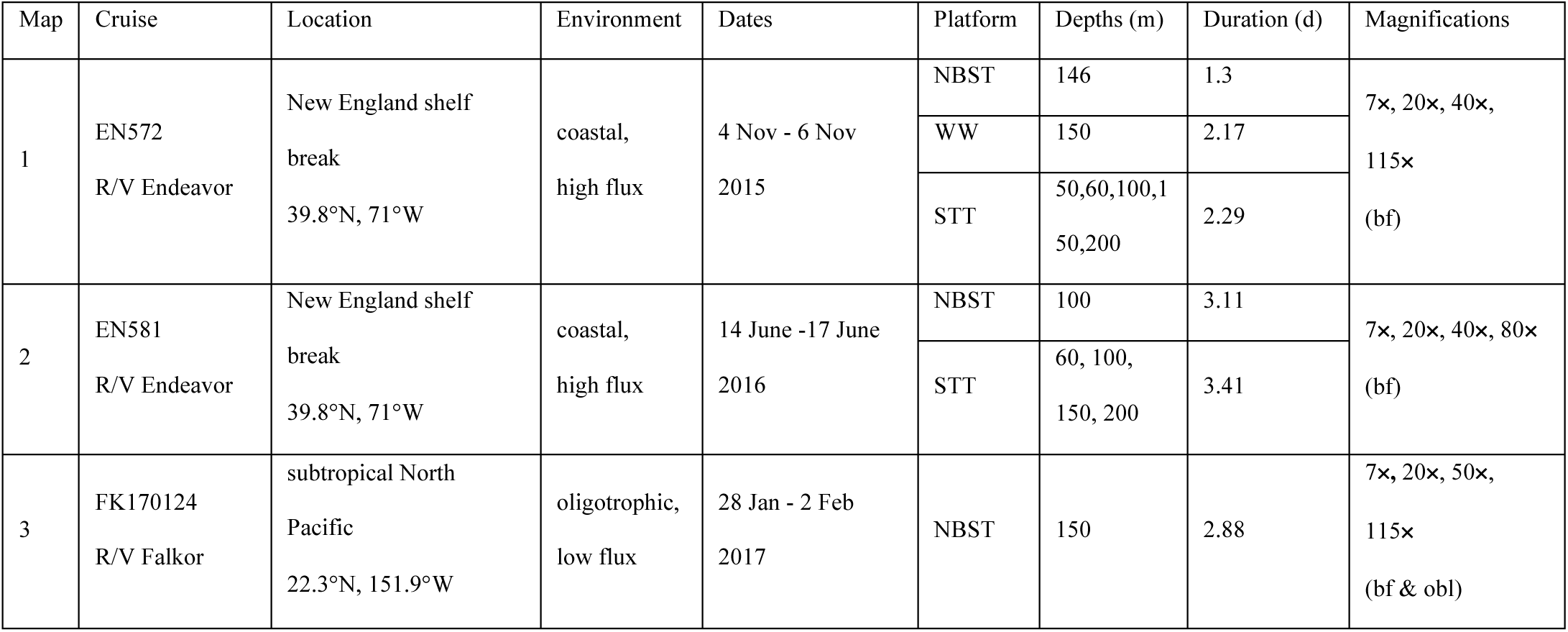
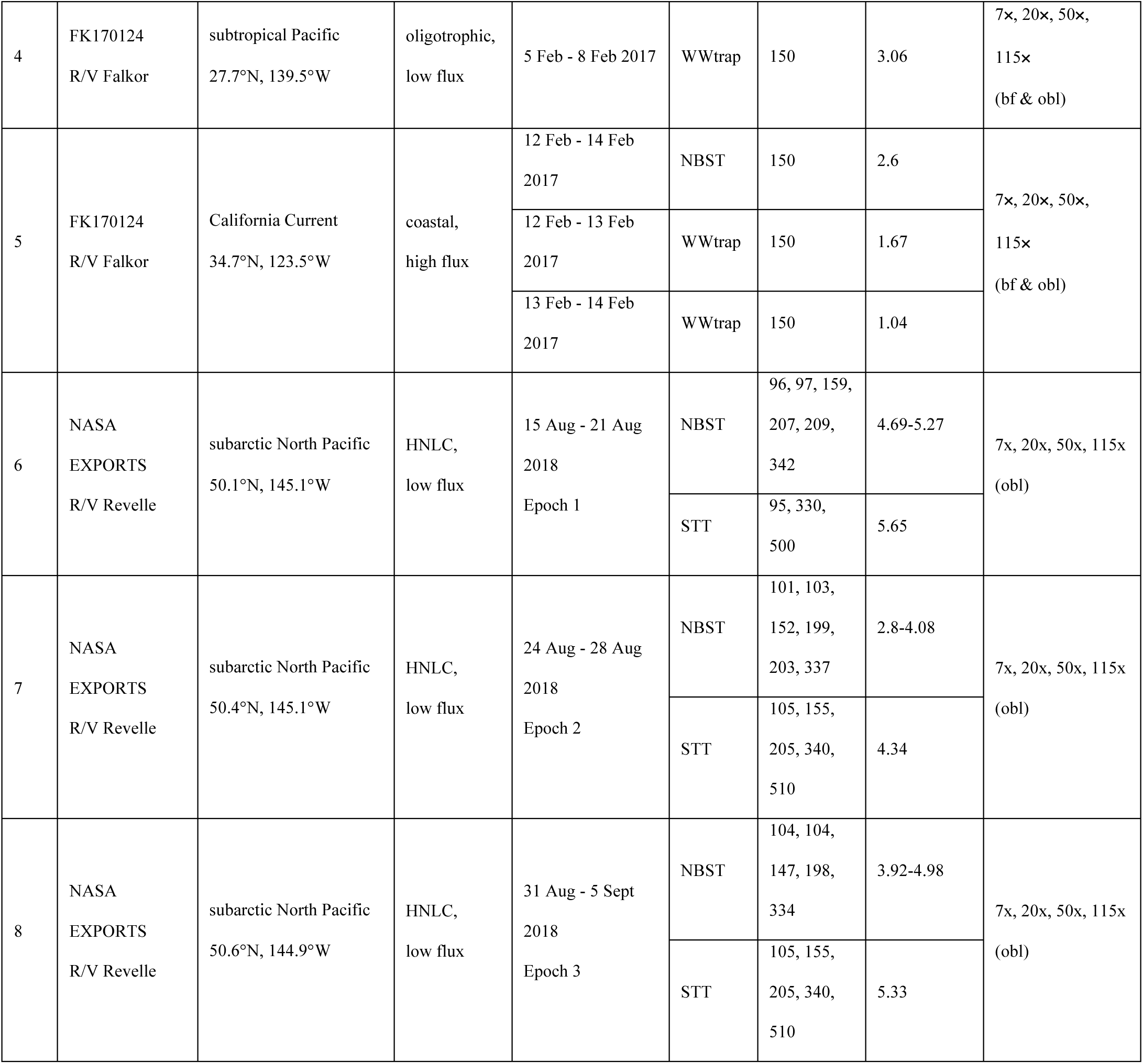
Summary of sampling locations labeled on map in Supplemental Figure 1, including cruise, deployment location, environment type, deployment dates, trap platforms, collection depths (meters), deployment duration (days), and imaging magnifications and illumination method. Environments were considered “high” flux if POC fluxes were > 2 mmol m^-2^ d^-1^. bf=brightfield, obl=oblique, HNLC=high nutrient low chlorophyll, NBST=neutrally-buoyant trap, STT=surface-tethered trap, WWtrap=Wirewalker trap.

| Map | Cruise | Location | Environment | Dates | Platform | Depths (m) | Duration (d) | Magnifications |
| --- | --- | --- | --- | --- | --- | --- | --- | --- |
| 1 | EN572<br>R/V Endeavor | New England shelf<br>break<br>39.8°N, 71°W | coastal,<br>high flux | 4 Nov - 6 Nov<br>2015 | NBST | 146 | 1.3 | 7×, 20×, 40×,<br>115×<br>(bf) |
|  |  |  |  |  | WW | 150 | 2.17 |  |
|  |  |  |  |  | STT | 50,60,100,150,200 | 2.29 |  |
| 2 | EN581<br>R/V Endeavor | New England shelf<br>break<br>39.8°N, 71°W | coastal,<br>high flux | 14 June -17 June<br>2016 | NBST | 100 | 3.11 | 7×, 20×, 40×, 80×<br>(bf) |
|  |  |  |  |  | STT | 60, 100, 150, 200 | 3.41 |  |
| 3 | FK170124<br>R/V Falkor | subtropical North<br>Pacific<br>22.3°N, 151.9°W | oligotrophic,<br>low flux | 28 Jan - 2 Feb<br>2017 | NBST | 150 | 2.88 | 7×, 20×, 50×,<br>115×<br>(bf & obl) |
| 4 | FK170124<br>R/V Falkor | subtropical Pacific<br>27.7°N, 139.5°W | oligotrophic,<br>low flux | 5 Feb - 8 Feb 2017 | WWtrap | 150 | 3.06 | 7x, 20x, 50x,<br>115x<br>(bf & obl) |
| 5 | FK170124<br>R/V Falkor | California Current<br>34.7°N, 123.5°W | coastal,<br>high flux | 12 Feb - 14 Feb<br>2017 | NBST | 150 | 2.6 | 7x, 20x, 50x,<br>115x<br>(bf & obl) |
|  |  |  |  | 12 Feb - 13 Feb<br>2017 | WWtrap | 150 | 1.67 |  |
|  |  |  |  | 13 Feb - 14 Feb<br>2017 | WWtrap | 150 | 1.04 |  |
| 6 | NASA<br>EXPORTS<br>R/V Revelle | subarctic North Pacific<br>50.1°N, 145.1°W | HNLC,<br>low flux | 15 Aug - 21 Aug<br>2018<br>Epoch 1 | NBST | 96, 97, 159,<br>207, 209,<br>342 | 4.69-5.27 | 7x, 20x, 50x, 115x<br>(obl) |
|  |  |  |  |  | STT | 95, 330,<br>500 | 5.65 |  |
| 7 | NASA<br>EXPORTS<br>R/V Revelle | subarctic North Pacific<br>50.4°N, 145.1°W | HNLC,<br>low flux | 24 Aug - 28 Aug<br>2018<br>Epoch 2 | NBST | 101, 103,<br>152, 199,<br>203, 337 | 2.8-4.08 | 7x, 20x, 50x, 115x<br>(obl) |
|  |  |  |  |  | STT | 105, 155,<br>205, 340,<br>510 | 4.34 |  |
| 8 | NASA<br>EXPORTS<br>R/V Revelle | subarctic North Pacific<br>50.6°N, 144.9°W | HNLC,<br>low flux | 31 Aug - 5 Sept<br>2018<br>Epoch 3 | NBST | 104, 104,<br>147, 198,<br>334 | 3.92-4.98 | 7x, 20x, 50x, 115x<br>(obl) |
|  |  |  |  |  | STT | 105, 155,<br>205, 340,<br>510 | 5.33 |  |

### Sediment trap sampling

To prepare tubes for deployment, seawater was collected using a CTD rosette or directly from the ship seawater flow through system, and filtered through a 1 *μ*m filter cartridge. Trap tubes were filled with filtered water overlying either 500 mL of 0.1% formalin brine (0.6 M NaCl, 0.1% formalin, 3 mM borate buffer) or a jar containing a polyacrylamide gel layer that was approximately 1 cm thick (Durkin et al. 2015). Identically prepared tubes were incubated in parallel onboard the ship to serve as process blanks. Upon recovery, collection tubes were allowed to settle for at least 1 h before the water overlying the gel was siphoned off. Jars containing polyacrylamide gel were removed from trap tubes and the remaining overlying water was carefully pipetted off the gel. Gels were stored at 4 °C and most were imaged within the following 2 days before being stored at −80 °C (see imaging methods below). Tubes containing formalin brine were drained through a 350 *μ*m or 335 *μ*m nylon mesh in order to separate and collect zooplankton that swam into the sample. These zooplankton “swimmers” were visually detected on the mesh using a dissecting microscope (Olympus SZX16) and removed using clean forceps. Non-zooplankton particles remaining on the mesh screens were rinsed back into the sample. On the EXPORTS and Falkor cruises, formalin-brine tubes were combined, and then split into 8 equal fractions using a custom built sample splitter (Lamborg et al. 2008). On the Endeavor cruises, the tubes from a given depth were combined into a single sample. Sample fractions were filtered onto precombusted, 25 mm glass fiber or quartz microfiber filters (Whatman GF/F or QMA) for particulate organic carbon quantification. Particulate organic carbon (POC) was determined using slightly different methodologies among research cruises and is described in detail in the Supplementary Methods. Briefly, POC was measured directly in some samples by combustion elemental analysis after fuming filters with concentrated acid to remove particulate inorganic carbon (PIC). In other samples, POC fluxes were calculated by separately measuring PIC and subtracting those values from the total particulate carbon measured by combustion elemental analysis (see Supplementary Methods).

### Gel trap sample imaging

Polyacrylamide gel layers were imaged on a dissecting microscope (Olympus SZX16) with either a Lumenera Infinity 2 (FK170124 and EXPORTS) or an Allied Vision Technologies StingRay (EN572 and EN581) camera attachment. The Lumenera camera resolved small particles better than the StingRay camera due to the greater pixel resolution. Particles collected in gel layers during EN572 and EN581 were imaged under brightfield illumination. Particles collected in gel layers during EXPORTS were imaged under oblique illumination. Particles collected in gel layers during FK170124 were imaged under both brightfield and oblique illumination, generating two separate sets of images for each sample and were used to determine whether illumination method affected particle detection and modeled POC fluxes. The POC fluxes modeled from brightfield illuminated samples were slightly lower than obliquely-lit samples due to lower detection of POC fluxes by small particle types and semi-translucent aggregates, though the uncertainty in total modeled fluxes often overlapped (Supplemental Figure 2). Data collected at the New England shelf break from brightfield illuminated images therefore most-likely underestimated fluxes by small and translucent particles.

All gel layers were imaged at 4 increasing magnifications, though the combination of magnifications varied by cruise (Table 1). Images were captured of non-overlapping fields of view spatially distributed across the gel area that excluded jar edges. At 7×, nearly every area of the gel surface was imaged, excluding areas near the jar edges. At 20×, a grid of images was randomly distributed across the entire gel surface. At 40× and 50×, images were collected across two orthogonal transects of the gel diameter. At 80× and 115×, images were collected across one transect of the gel diameter. At magnifications greater than 7×, multiple focal planes within a field of view were imaged to capture particles embedded in different depths of the gel layer and varied between two and nine focal planes. The number of focal planes imaged was consistent across all fields of view for a given magnification but varied across cruises due to variation in gel thickness and the vertical distribution of particles in the gel. All images from the Falkor and Endeavor cruises are available at www.bco-dmo.org/project/675296 (last accessed May 29, 2020). Images from the EXPORTS cruise are available at seabass.gsfc.nasa.gov/archive/MLML/durkin/EXPORTS/exportsnp/associated (last accessed May 29, 2020).

Some of the gel layers collected during the EXPORTS cruise were imaged after being stored at −80 °C and thawed. To determine whether measured particle properties changed after gel layers were frozen, samples collected during FK170124 were thawed after being stored for approximately 1 year at −80 °C and imaged again under both brightfield and oblique illumination. The POC fluxes modeled from obliquely lit particles did not differ after gels had been frozen (Supplemental Figure 3) but were consistently lower when imaged under brightfield illumination. Freezing samples prior to imaging should not affect inter-sample comparisons when imaged under oblique illumination.

### Image processing and particle detection

Particles in gel images were quantified with an image processing protocol created using functions available in python’s Sci-Kit Image (Python code available at github.com/cadurkin, accessed May 29, 2020 and in Supplemental File 1). The protocol (Fig. 1) is similar to that reported by Huffard et al. (2020) and described in detail in the Supplemental Methods. Briefly, particles present in each micrograph were detected through a series of image transformation steps that maximize the detection of entire particle areas while minimizing the detection of imaging noise. The background was removed, a brightness threshold was applied, and an edge-detection kernel transformation was used to identify the in-focus particles (Fig. 1a). Duplicate particles detected in multiple focal planes were removed and the remaining particles were counted and measured (Fig. 1b).

### Assigning particle image identities

Particles were categorized into 9 different sinking particle classes (Fig. 2, Table 1). Identities were assigned by manually identifying particle images. Aggregates were defined as loosely packed detritus with irregular edges. Dense detritus was defined as densely packed amorphous material, often brown or golden in color. Large, loose pellets were similar to dense detritus but were also elongated like a fecal pellet. We presume that the main zooplankton source of these pellets in our samples was pyrosomes, a type of colonial pelagic tunicate. This assumption is based on our observations of pyrosomes in the California Current, where we observed most of these pellets. Pyrosomes were unintentionally caught on top of the trap baffles and in the gel layers, generated intense bioluminescence at night, and were observed in situ in high abundance several weeks later during a separate study with a remotely operated vehicle (Durkin, personal observation). Long fecal pellets were defined as long, thin, cylindrical fecal pellets with a smooth edge, such as the chitin-encased pellets produced by euphausiids. Short fecal pellets were also smooth-edged pellets but with an oval or ellipsoid shape, such as those produced by larvaceans. Mini pellets were defined as small (usually <100 *μ*m ESD), spheres such as those produced by rhizarians and microzooplankton (Gowing and Silver 1985). A small number of individual organisms considered to be passively sinking were also detected, including Rhizarians, primarily Phaeodaria, and various phytoplankton, usually diatoms. Pteropods, copepods, amphipods, foraminifera, and other zooplankton that probably swam into the gel and human-produced fibers were also detected but not considered passive flux and not included in the categories of sinking particles. Unidentifiable objects were also detected and were likely caused by out-of-focus particles, shadows, smudges, or noise detected by the image processing steps that are sensitive to the particular thresholds used. The percent of particle candidates that were unidentifiable increased with decreasing size; >50% of particle candidates smaller than 40 *μ*m fell into this category whereas ∼20% of all particle candidates larger than 40 *μ*m could not be identified. The increase in imaging noise or contamination in smaller particle sizes classes is consistent with the trends in background noise detected in gel process blanks (Durkin et al. 2015).

**Figure 2.**
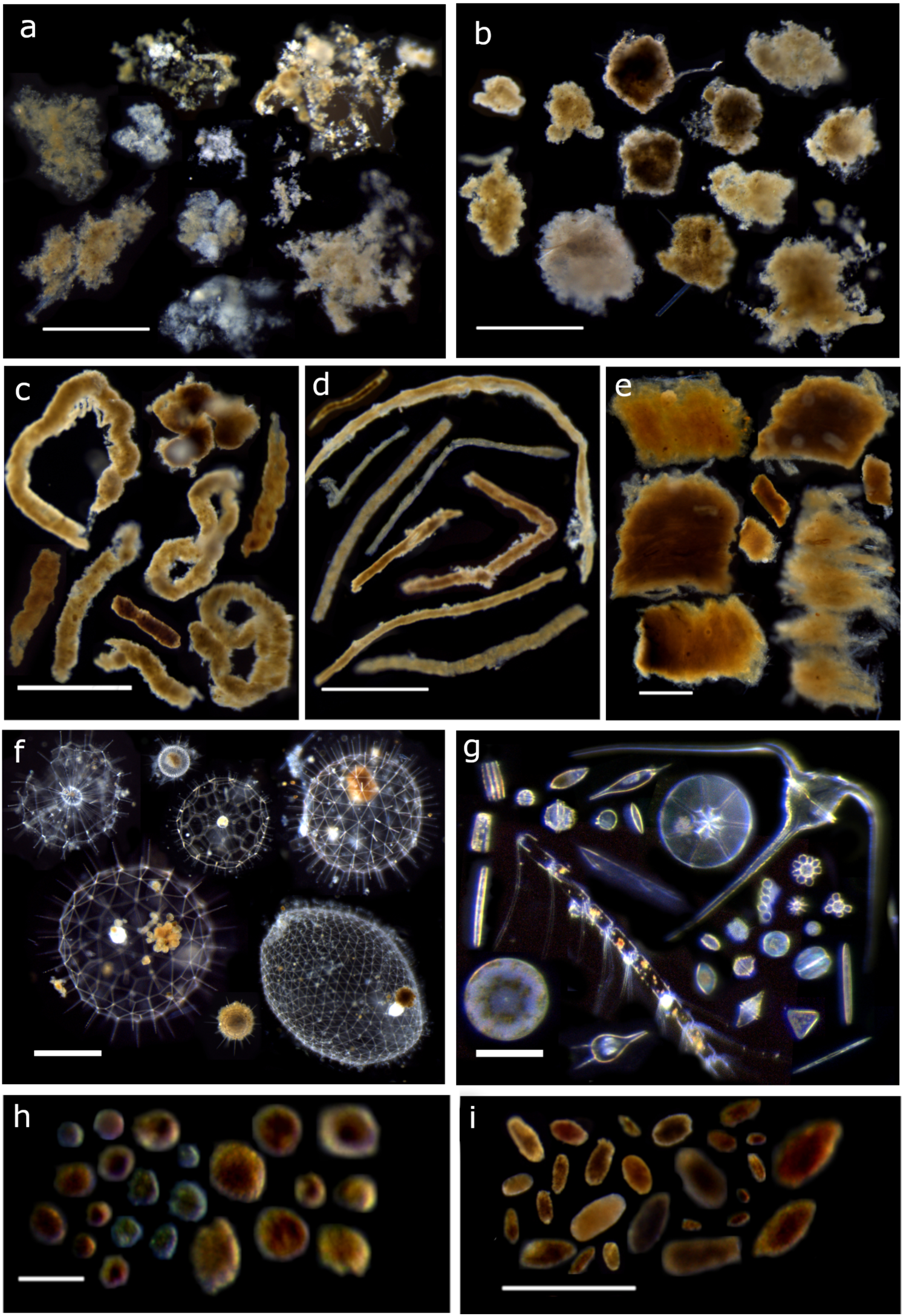
Example micrograph collages of each sinking particle classification, including a) aggregates, b) dense detritus, c) large loose fecal pellets, d) long cylindrical fecal pellets, e) salp fecal pellets, f) Rhizaria, primarily Phaeodarians, g) phytoplankton cells, h) mini pellets, i) short (ellipsoid or oval) fecal pellets. Scale bars in panels g and h: 100 *μ*m, scale bars in panels a-f and i: 1000 *μ*m. Examples include particles collected across multiple locations.

### Calculating particle fluxes

Only identifiable, sinking particle classes from the dataset were considered in calculations of sinking flux. For each imaging magnification, particles were grouped by their equivalent spherical diameter (ESD) into logarithmically-scaled size bins with edges at (2^#^)|^12^ *μ*m. To calculate number fluxes at each magnification, the number of particles counted in each size bin was divided by the total imaged surface area of the gel and the trap collection time. Uncertainty of the number flux was estimated by applying these same calculations to the counting uncertainty (square root of the number of particles counted, ie. Poisson distribution). Each imaging magnification had an optimal size range for resolving particles and these ranges were combined from each magnification to generate the combined number flux (Fig. 1c, Durkin et al. 2015). These size ranges differed slightly between images collected during both cruises at the New England shelf break using the StingRay camera versus the images collected during the Pacific Ocean cruises using the Lumenera camera.

### Modeling carbon fluxes

The carbon flux by each particle type was determined by modeling particle volumes, calculating the carbon per unit volume, and multiplying the carbon per particle by the measured particle number fluxes in each size category. Equations used to model the volume of each particle type are listed in Table 2. To estimate the volume of non-spherical fecal pellets based on the automated image measurements, a relationship between pellet width and ESD was established as follows. First, widths of a subset of pellets collected at all sampling locations were manually measured using tools in ImageJ (Schneider et al. 2012) and included 1415 long fecal pellets, 563 large loose fecal pellets, 596 short fecal pellets, and 186 salp fecal pellets (Supplemental Fig. 4). The biological basis for a linear relationship between width and ESD is that larger animals can produce correspondingly wider pellets with a larger total ESD. This direct linear relationship can be used to estimate the width of short fecal pellets and salp fecal pellets. The width:ESD of short fecal pellets was 0.54, with a normalized root mean squared difference (NRMSD) of 2%. The width:ESD of salp pellets was 0.63 (NRMSD = 6%). This linear relationship is not valid when long pellets fragment into smaller pieces that still have a large width, as occurs with the long fecal pellets and the large loose fecal pellets (Supplemental Fig. 4). The relationship between width (w, in *μ*m) and ESD of these pellet types was instead described by a hyperbolic equation.

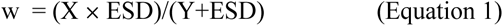

**Table 2.**
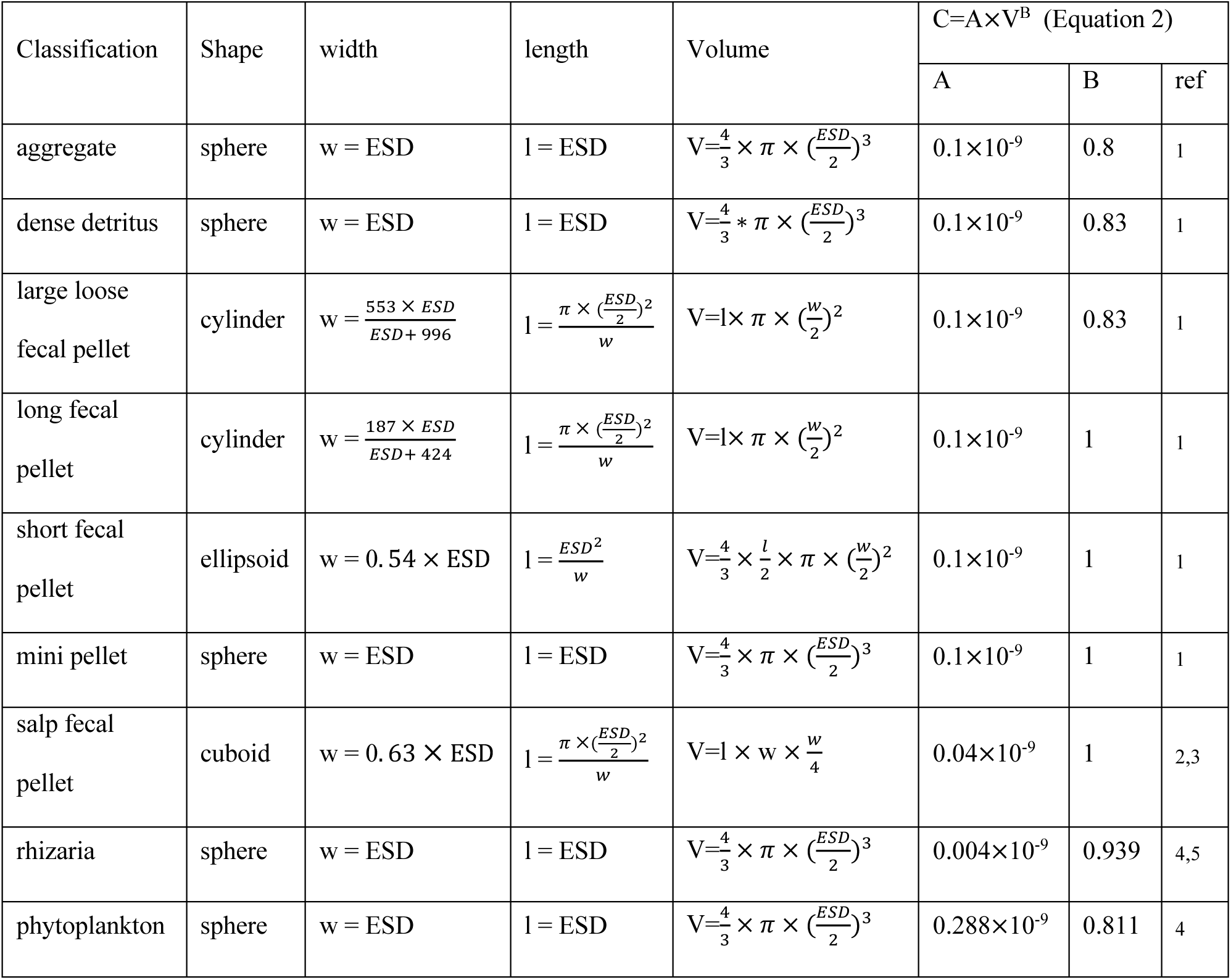
Assumptions and equations used to model the particulate organic carbon content of each particle type. Parameters for equation 2 were used from various reference studies: ref 1: this study, 2: Silver and Bruland (1981), 3: Iversen et al. (2017), 4: Menden-Deuer and Lessard (2000), 5: Stukel et al. (2018). ESD=equivalent spherical diameter.

The best-fit parameters of X and Y were 187 *μ*m and 424 *μ*m for long fecal pellets (NRMSD=7%) and 553 *μ*m and 996 *μ*m for large loose fecal pellets (NRMSD=9%). Pellet lengths were calculated for each pellet shape given the modeled widths and measured 2-dimensional area (Table 2). Length (l) and width (w) were used to calculate the volume of each pellet in a given size category. Salp fecal pellets were modeled as cuboids where pellet depth was assumed to be one quarter of the width (Silver and Bruland 1981). Particle volumes were converted to carbon units using published relationships and best-fit parameters to Equation 2

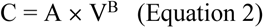

where C is the carbon per particle (mg), A is a scaling coefficient (mg *μ*m^-3^), V is particle volume (μm^-3^) and B is a unitless exponent parameter.

The *optimize.minimize* function in python was used to find the values of A and B for particle categories in the first six rows of Table 2 that gave the best fit to log transformed, bulk carbon fluxes measured in trap tubes. In the fitting process, the same value of A was assumed for all six of these particle categories and B=1 was assumed for all compact fecal pellet categories (long, short, mini). Shared B values were assumed for dense detritus and large, loose fecal pellet and a separate B value was assumed for aggregates. Literature values of A and B were applied to salp fecal pellets, rhizaria, and phytoplankton (Table 2). In total, three parameters were fit to 13 observations from trap platforms that carried both a polyacrylamide gel layer and poisoned brine (Fig. 3), and included samples from location 1,2,3 and 5 in Table 1. Observations made during the EXPORTS cruise in the subarctic N. Pacific were not included in fitting parameters because gel images were used to correct POC fluxes and therefore were not an independent measurement (Estapa et al. in review). The gel particle sample collected at one subtropical station (location 4, Table 1) did not have a corresponding bulk POC flux measurement. Only one of the three gel particle samples collected in the California Current had a corresponding bulk POC flux sample collected on the same platform. The standard error of model parameters was estimated by the square root of the inverse Hessian matrix.

**Figure 3.**
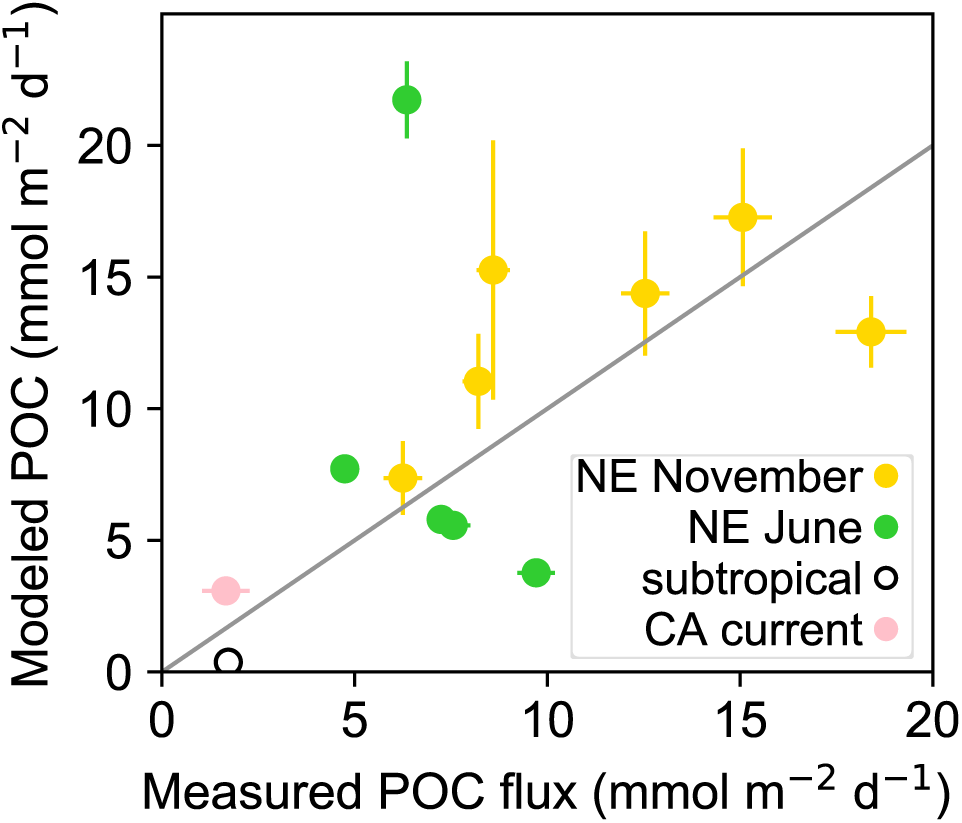
Particulate organic carbon (POC) fluxes modeled from particle images versus measured from formalin-preserved bulk collector tubes. Black line equals 1:1 relationship. Vertical error bars represent modeled uncertainty in POC flux calculated from the propagated counting uncertainties of particles (Poisson distribution). Horizontal error bars represent analytical uncertainty of combined triplicate tubes for New England (NE) shelf deployments and the standard deviation among splits for subtropical Pacific and California (CA) current deployments.

The best-fit value of A for the six modeled particle categories (0.1×10^-9^ ± 0.04×10^-9^ mg *μ*m^-3^) falls in the middle of the range of reported carbon densities for fecal pellets (González et al. 1994; Lundsgaard and Olesen 1997; Urban-Rich et al. 1998). The values of B less than 1 represent the effect of packaging differences among particle types, with relatively lower carbon density as particle sizes increase as found by Alldredge (1998). The aggregate B value was 0.80 ± 0.05, and dense detritus and large loose fecal pellet B value was 0.83 ± 0.11. Values of A and B were different than those calculated for aggregates by Alldredge (1998) who measured particles larger than 1 mm^3^ while nearly all of aggregates observed in this study were smaller. Our parameters predict the same aggregate POC contents as Alldredge’s parameters for aggregates with an ESD of 709 *μ*m. Particles smaller than this size are predicted to have a lower POC content than would be predicted by Alldredge’s parameters. The best fit parameters for non-aggregate particle types was not sensitive to whether we used Alldredge’s aggregate parameters or instead fitted our own. Uncertainty of modeled POC fluxes for a given size bin and particle class were calculated by multiplying the modeled carbon contents by the calculated number flux uncertainties. The uncertainties within each size bin were propagated to estimate the uncertainty of the total modeled POC flux by each particle type integrated across all sizes. All particle counts, fluxes, modeled volumes, and modeled POC fluxes across size bins are presented in Supplemental File 2, including counts of non-sinking particle classes.

### Statistical analyses

A principal component analysis (PCA) was performed to identify groups of samples with similar relative particle compositions. The modeled POC flux of each particle type in a sample was normalized to the total modeled carbon flux in the sample. The covariance matrix of this normalized data was calculated using python function *numpy.cov*. The *scipy.linalg.eig* function was used to calculate eigenvalues and vectors from which component loadings and scores were determined.

Flux attenuation was assessed for each particle type at locations where traps were deployed at multiple depths. Flux attenuation slopes of each particle type from all cruise data was calculated using the Martin (1987) power law equation:

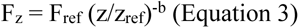

where carbon flux (F_z_) of any particle type at depth (z) is determined by the attenuation coefficient (b) and the flux at the reference depth (F_ref_). The reference depth was considered to be the depth at which photosynthetically active radiation (PAR) was at 1% of surface values. This depth was 78 m in the subarctic North Pacific and at 58 m and 43 m at the New England shelf break in November and June, respectively. The least-squares, best-fit values for b and F_ref_ describing log-transformed modeled carbon fluxes were calculated using python’s *optimize.minimize* function. The choice of reference depth only affects the best fit value of F_ref_, and does not influence the best fit value of b. To calculate the uncertainty of these parameter estimates, we generated 200 simulated datasets of modeled POC flux, distributed randomly within the normal distribution of the measurement uncertainty, which was based on the number of particles counted. The mean and standard deviation of modeled parameters were calculated from all simulations in which the minimization function converged on model parameters. Attenuation curves were fit to the data only if the standard deviation of modeled parameters (our measure of uncertainty) was less than 50% of the parameter value. If the minimization function never converged on model parameters or the standard deviation of parameter values was greater than 50%, the data were considered to be inconsistent with a power law flux attenuation model.

## Results

### Reproducibility and bias across samples

Modeled POC fluxes reported here may be affected by stochastic collection of particles in the trap tubes, small-scale spatial variation in particle fluxes among trap platforms, and collection biases of the trap platforms that collected the fluxes. To assess the influence of stochastic particle collection, triplicate gel layers were deployed on the same Wirewalker trap platform during the November cruise to the New England shelf break (Supplemental Fig. 5a). Modeled POC fluxes among these three samples were similar, with small variation among replicates (maximum difference of 2.2 mmol C m^-2^ d^-1^, or 35% of the mean flux). The effect of small-scale spatial variability on modeled POC flux was assessed across duplicate neutrally-buoyant trap platforms deployed at the same time and depth in the subarctic North Pacific (Supplemental Fig. 5b). On 2 of 5 occasions the uncertainties in modeled POC fluxes did not overlap and these differences (1.1-2.2 mmol C m^-2^ d^-1^, or 43-113% of mean flux) were slightly higher than differences among replicate tubes on the same platform (Supplemental Fig. 5a compared to 5b). To determine whether modeled POC fluxes were affected by trap platform type, replicate traps of different platform types were co-deployed on 13 occasions (Supplemental Fig. 5c, 5d). The modeled POC flux collected by the Wirewalker trap did not differ from the flux collected by the neutrally-buoyant trap on the two occasions when they were co-deployed, suggesting that this unconventional trap design functioned as intended. The modeled POC flux collected by the surface-tethered trap was greater than that collected by the neutrally-buoyant trap on 8 out of 12 occasions and was never lower than the neutrally-buoyant trap. This apparent collection bias is consistent with measured differences in POC fluxes collected by these two platform types (Estapa et al. in review). Differences between the modeled POC fluxes collected by the surface-tethered and neutrally-buoyant traps were comparable to differences caused by small scale spatial variation (Supplemental Fig 5b) (0.17-7.9 mmol C m^-2^ d^-1^, or 31-114% of mean flux). Overall, the degree of variability in modeled POC flux among samples and platforms was consistent with previous sediment trap studies addressing variability in bulk chemical fluxes (Owens et al. 2013; Baker et al. 2020).

### Flux composition patterns across sites

Particle export was similar at two locations sampled in the North Pacific subtropical gyre in February 2017. All sinking particles were smaller than 1000 *μ*m and primarily composed of aggregates, dense detritus, and short fecal pellets (Fig. 4). Particles smaller than 100 *μ*m were largely composed of mini pellets (7% and 9% of total modeled POC flux at locations 3 and 4). In the California Current, particles smaller than 1000 *μ*m were similar in composition to those in the subtropical locations (aggregates, dense detritus, and short fecal pellets, mini pellets). Modeled POC fluxes in the California Current were also composed of particles larger than 1000 *μ*m including long fecal pellets (35±6% of total flux) and large, loose fecal pellets likely produced by pyrosomes (10±2% of total flux) (Fig. 4). Phaeodarian rhizaria also accounted for 18±14% of the POC flux by particles >1000 *μ*m (3±3% of the total modeled POC flux).

**Figure 4.**
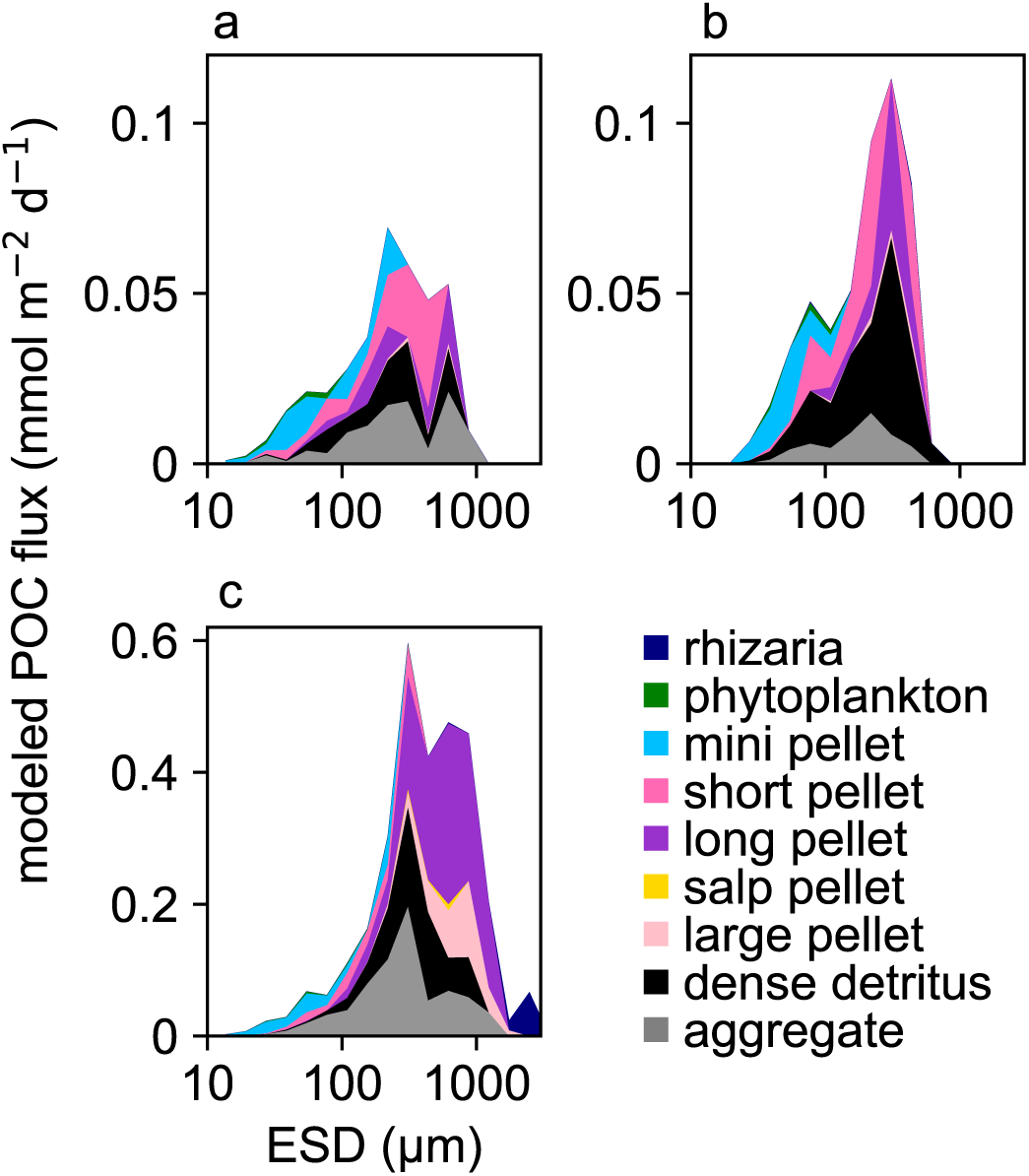
Differences in wintertime sinking particle composition in the North Pacific at a&b) two subtropical locations (locations 3 and 4, Table 1) and c) the California Current at a depth of 150 m in February 2017. Colors indicate the contribution of each particle type to the modeled particulate organic carbon (POC) flux in each particle size bin measured as equivalent spherical diameter (ESD). The mean of all three samples collected concurrently in the California current are shown.

Carbon flux at the New England shelf break in November 2015 was dominated by particles larger than 1000 *μ*m at all depths (50 m - 200 m; 63 ± 14%), largely composed of phytodetrital aggregates (36 ± 11%) and dense detritus (17 ± 7%) that visibly contained large diatom cells (e.g. *Coscinodiscu*s; Fig. 5a). Salp fecal pellets and long fecal pellets composed most of the remaining POC flux (14 ± 11% and 27 ± 6%, respectively). In November, modeled fluxes attenuated with depth similar to measured POC fluxes whereas in June the modeled POC fluxes did not match the measured POC fluxes profile as well. June particle fluxes at this location (Fig. 5b) were very different from November, with long fecal pellets exporting nearly all sinking carbon at all depths (60-200 m; 83±8%) and attenuated rapidly in the upper 100 m.

**Figure 5.**
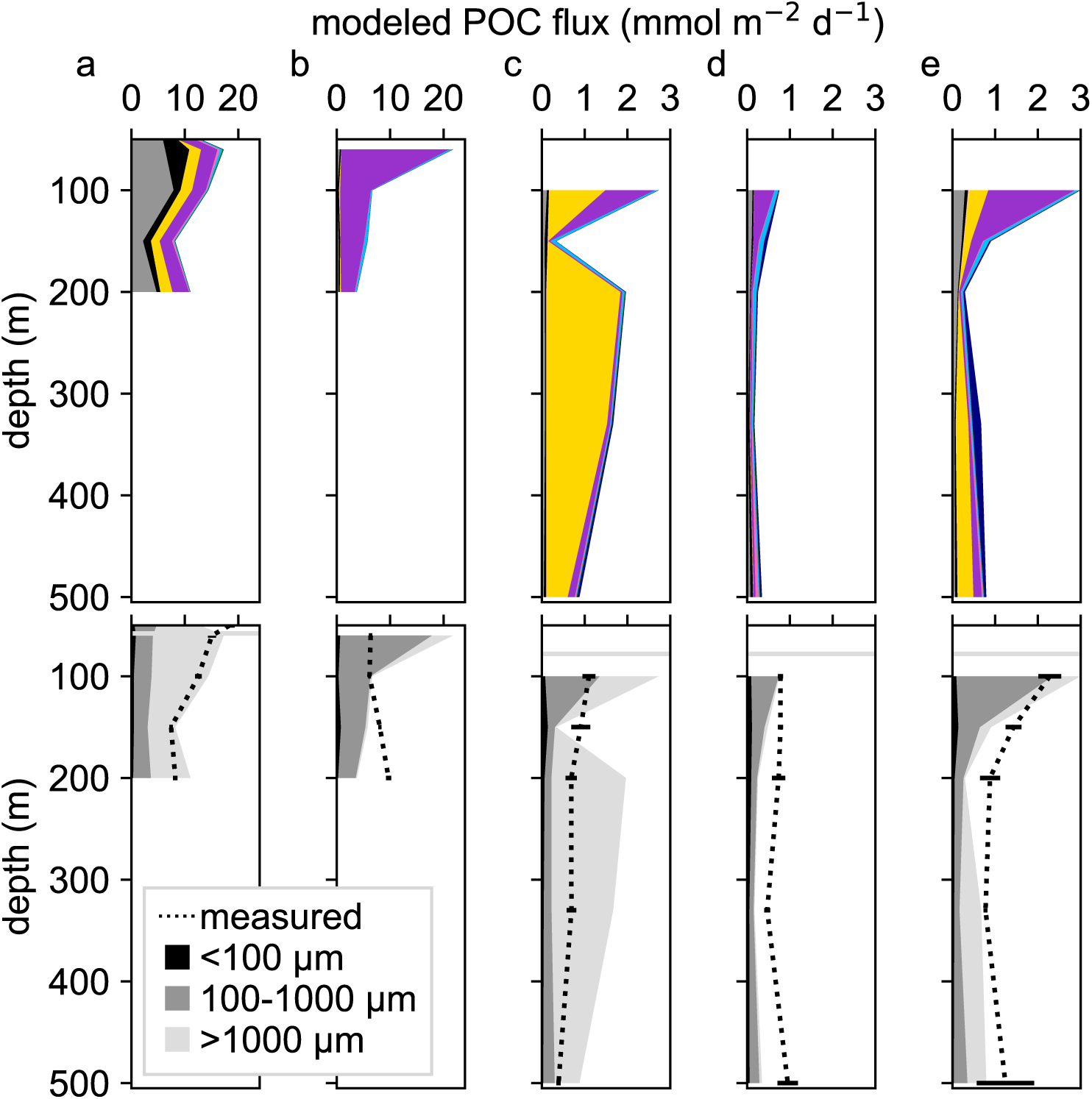
Depth profiles of mean modeled particulate organic carbon (POC) flux a) at the New England shelf break in November 2015 and b) June 2016 and in the subarctic North Pacific during successive sampling periods c) epoch 1, d) epoch 2, and e) epoch 3. When replicate trap platforms were deployed at the same depth horizon values indicate the mean of modeled fluxes (see Table 1). Top figure in each panel shows modeled POC flux by each particle type, with colors corresponding to the key in Figure 4. Bottom figure in each panel shows mean modeled POC flux by three size categories of particles collected in all traps. Dashed line indicates average measured POC fluxes collected in formalin-poisoned trap tubes. Error bars indicate the analytical uncertainty of three combined trap tubes. Horizontal grey line indicates depth of 1% surface photosynthetically active radiation.

Modeled carbon flux in the subarctic North Pacific was an order of magnitude lower at all observed depths (95 - 510 m), and matched the scale and general depth profile of measured fluxes (Estapa et al. in review) even though carbon modeling parameters were not fit from this location (Fig 5c-e). Particle composition varied drastically with depth and across the three deployment periods. Long fecal pellets were a major component of carbon export out of the euphotic zone and attenuated rapidly with depth. In spite of their small size (<100 *μ*m), numerically abundant mini pellets contributed between 7% and 36% of the total carbon flux (mean across depths: 17 ± 9%). Salp fecal pellet fluxes were numerically rare (usually 1 or 2 pellets observed per sample and never more than 7), but had an outsized influence on the magnitude and of carbon flux and varied across the sampling periods.

### Relationship of particle compositions among samples

The composition of sinking particles was highly variable across coastal, open ocean, temperate, and subtropical environments (n=49 samples). To identify samples with similar particle compositions, a PCA was performed on the fractional composition of each particle type within each sample (flux of each particle type normalized to total of all types in a sample; Fig. 6). Variability in relative particle composition among samples was primarily driven by the contribution of three particle classes to carbon flux: aggregates, long fecal pellet, and salp fecal pellets (Fig. 6a). The first principal component accounted for 51% of the variance, and was dominated by an inverse relationship between relative salp fecal pellet flux and relative long fecal pellet flux. The second principal component accounted for 31% of the variance, was nearly parallel to increases in relative aggregate flux, and opposed to the relative fluxes of long fecal pellet flux and salp fecal pellet flux. Samples appeared to cluster along the directions of these three dominant particle classes and often separated by location or season (Fig. 6b). An exception to this pattern was found in the samples collected in the subarctic North Pacific (n=30) over a period of one month where fluxes were associated with all three of the major particle types (Fig. 6c). At this location, the dominant particle type varied with depth: long fecal pellets were exported just below the euphotic zone (∼100 m) while aggregates and salp fecal pellets contributed to most of the carbon export as depth increased.

**Figure 6.**
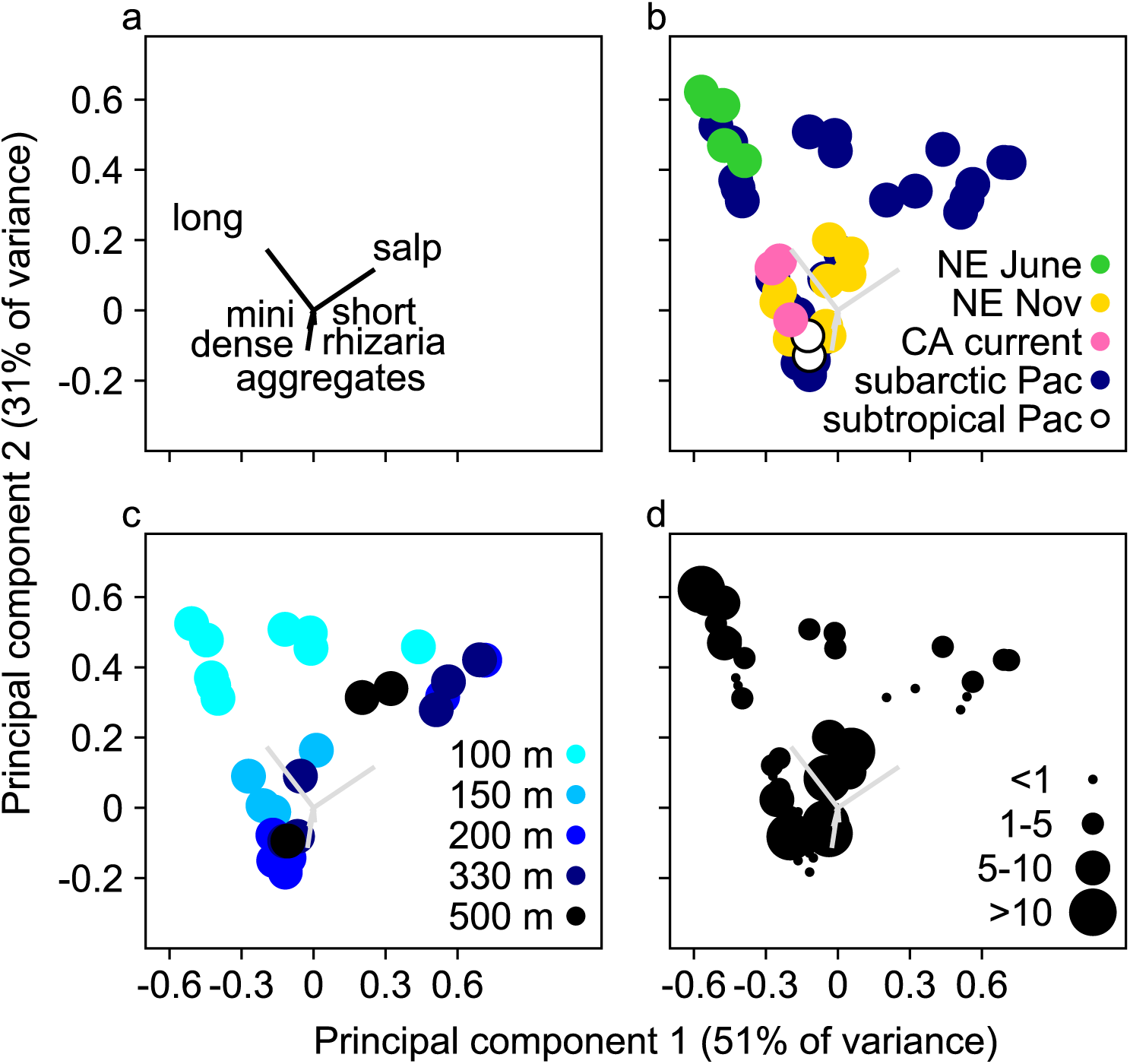
Principal component analysis of relative particle composition covariance from nine particle categories in all 49 samples. Carbon fluxes in each sample were normalized to total flux by all particle types prior to this analysis of the covariance matrix. a) Nine component vectors with seven labeled. Particles include various fecal pellet types (long, mini, salp, short), detritus (aggregates and dense), and individual cells such as rhizaria. Phytoplankton and large loose fecal pellet carbon flux vectors are near the origin and are not labeled. b) Variation in particle composition across locations and samples. Colors indicate sample location. c) Variation in particle composition in the subarctic North Pacific samples only. Colors indicate approximate sampling depth. d) Variation in particle composition among all sampling locations with relation to total modeled POC fluxes. Circles sizes indicate magnitude of total modeled POC fluxes in mmol C m^-2^ d^-1^. Samples were collected in the subarctic North Pacific (Pac), the New England (NE) shelf break, and the California (CA) current.

### Efficiency of carbon export among particle types

One, perhaps surprising, result to come from this analysis is that the magnitude of the modeled flux was not related to the particle type that primarily composed the fluxes. Each of the dominant particle types (long fecal pellets, aggregates, or salp fecal pellets) was associated with both high and low magnitude total carbon flux (Fig. 6d). Instead, each particle type had a different effect on the rate at which carbon attenuated with depth (Fig 7, Supplementary Table 2). Across a spectrum of flux environments and seasons, specific particle types shared similar attenuation profiles across locations and sampling periods. Aggregate fluxes out of the euphotic zone attenuated moderately with depth (b = -0.8±0.3). The attenuation slope of aggregates was poorly constrained (>70% uncertainty) when total aggregate fluxes were low in June at the New England shelf break, and was not included in this assessment. Long fecal pellet fluxes out of the euphotic zone attenuated more rapidly with depth (b=-1.8±0.3). The attenuation slope of long fecal pellets was poorly constrained at the New England shelf break in November (>90% uncertainty), and these samples were not considered in the assessment. Mini pellets also attenuated with depth (b=-1±0.1). The attenuation slope of mini pellets collected at the New England shelf break in June was highly uncertain (>70%) and not considered further. The other particles that contributed substantially to POC flux (salp fecal pellets, dense detritus, short fecal pellets) had depth profiles of flux inconsistent with exponential attenuation and, in some cases, fluxes may have increased with depth. (Fig. 7).

**Figure 7.**
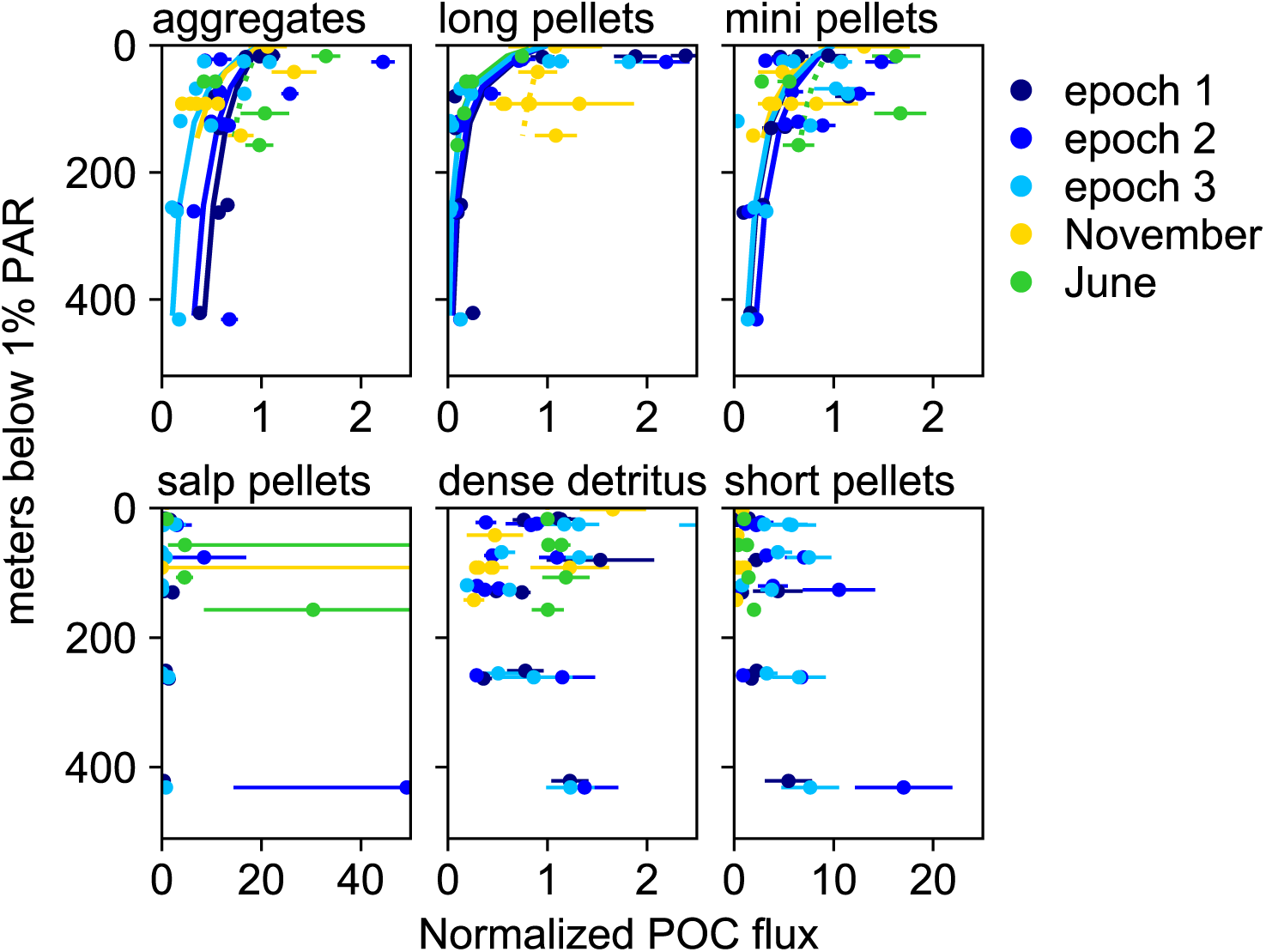
Changes in modeled particulate organic carbon (POC) flux with depth by major particle types identified by the principal component analysis (PCA). Large loose fecal pellets, phytoplankton and rhizaria fluxes not analyzed here because they were numerically rare at these locations and/or contributed little to the modeled POC flux. Fluxes were normalized to the modeled flux at the 1% photosynthetically active radiation (PAR) light level (top three panels). Particle types for which flux attenuation curves could not be modeled were normalized to the average flux of the shallowest traps (bottom three panels). Note changes in scale of the x-axes. Colors indicate sampling location where multiple depths were observed, including the subarctic North Pacific during all three sampling epochs, and at the New England shelf break in November 2015 and June 2016. Error bars represent the uncertainty of modeled POC flux calculated from the propagated uncertainty of particle counts (Poisson distribution). Solid lines indicate the slope of attenuation if uncertainties of this estimate were ≤ 50%. Dashed lines in top three panels indicate the slopes that did not fit this criterion.

### Contribution of small particles to carbon export

Particles with an ESD smaller than 100 *μ*m accounted for between 1% and 46% of modeled total POC fluxes (Fig. 8). In samples with relatively high total POC fluxes (2-22 mmol C m^-2^ d^-1^), the percent contribution by small particles was 5±3%. At locations where POC flux was lower than 2 mmol C m^-2^ d^-1^, the percent contribution of small particles was higher on average (19±13%).

**Figure 8.**
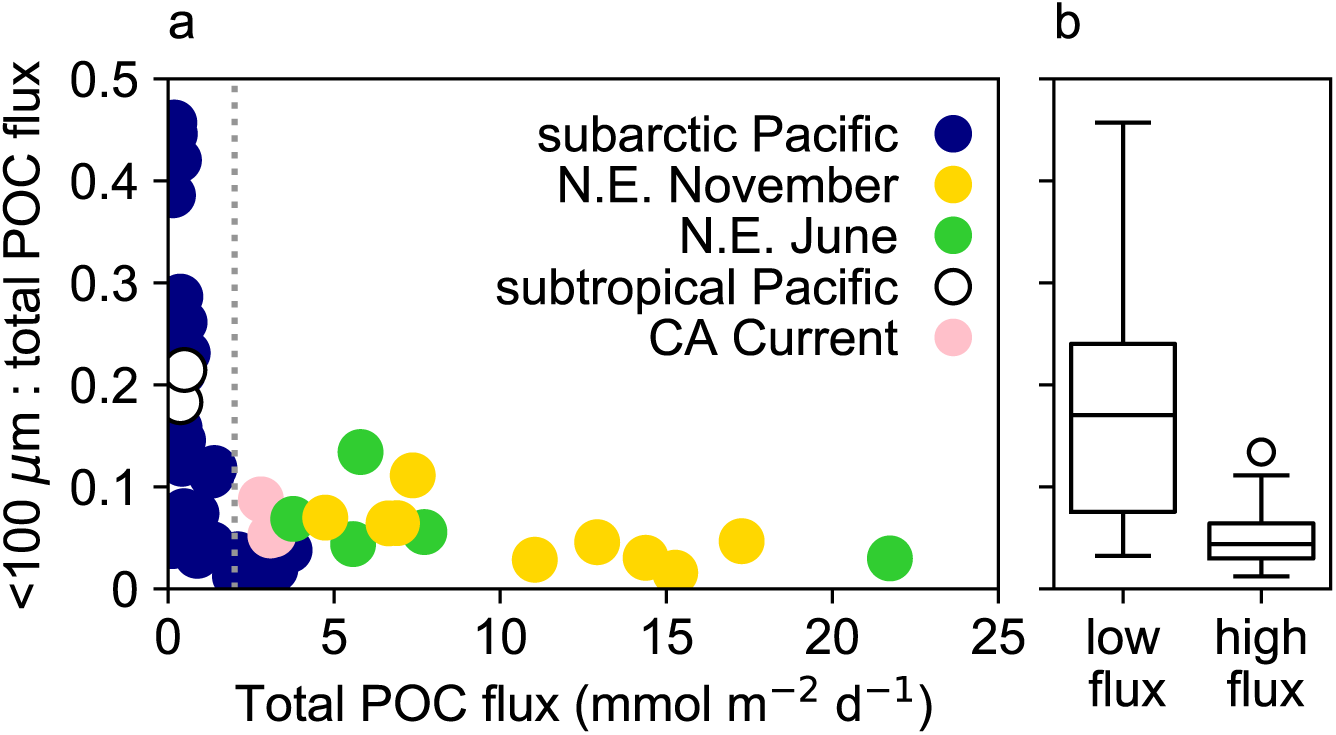
The contribution of small particles to total POC flux. a) Fraction of modeled particulate organic carbon (POC) flux by small (< 100 *μ*m) particles versus the total modeled POC flux. Vertical dashed line separates samples with relatively low (<2 mmol C m^-2^ d^-1^) and high modeled POC flux. b) Boxplot distribution of the fraction of POC flux by small particles in samples with low and high modeled POC flux. Horizontal center line indicates the mean, boxes extend to the first quartile, whiskers indicate the range of quartiles, and points indicate data outliers. NE = New England, CA = California.

## Discussion

We quantified the particle types responsible for gravitational carbon export across a range of ecosystems and seasons through detailed visual observations and measurements. These snapshots of carbon export indicated that the relative contribution of aggregates, long fecal pellets, and salp fecal pellets had the largest effect on the variability of POC composition among the 49 samples. Among the nine particle types that represent different carbon flux pathways, no single type explained the high or low flux magnitudes observed across these disparate environments. Instead, each particle type showed a different depth attenuation pattern. We suggest that identification of particles through imagery can reasonably improve the prediction of the magnitude of carbon flux and its potential to attenuate with depth.

Our observations provide momentary snapshots of the biological pump across very different ecosystems. Though we did not capture the full range of conditions that occur at any one location, some general patterns still emerge related to 1) the effect of trophic transformation of POC on flux attenuation, 2) the influence of episodic fluxes generated by gelatinous zooplankton, and 3) carbon export by small particles.

The transfer of carbon to depth appeared to be affected by the degree to which sinking particles were transformed by zooplankton grazers and microbes. Particles collected in the subtropical Pacific were primarily detrital (aggregates, dense detritus) and were likely the most trophicly-processed particles among all the locations sampled in this study. Based on the depth attenuation patterns observed at other locations, the abundant small particle types observed in this environment (mini pellets, short fecal pellets, dense detritus) likely attenuated little with depth. These findings are consistent with previous studies of the subtropical gyre that reported low export but high transfer efficiency (Buesseler and Boyd 2009). At other locations where sinking particles were primarily generated by mesozooplankton, the effect on flux attenuation was very different. The large and relatively fast-sinking pellets collected in the California Current, the New England shelf break in June, and in the subarctic North Pacific were likely produced by euphausiids, copepods, and amphipods and may have provided a more direct export pathway out of the surface ocean compared to the subtropical sampling locations. However, these long fecal pellets also attenuated very rapidly with depth (b=1.8±0.3), consistent with previous studies that measured high attenuation of crustacean fecal pellets in the upper water column (Alldredge et al. 1987; Urban-Rich et al. 1999). In the subarctic North Pacific, long fecal pellets sinking in the upper 100 m of the mesopelagic were likely fragmented and consumed by mesopelagic zooplankton, respired by bacteria, and/or transferred to other sinking particle classes (e.g. dense detritus, aggregates). The food supplied to mesopelagic organisms by fragmentation of long fecal pellets may ultimately be repackaged into other sinking pellet types (e.g. short pellets produced by larvaceans). Based on these observations, we hypothesize that when mesozooplankton fecal pellets drive fluxes out of the surface ocean, particle transformations in the mesopelagic will be relatively dynamic due to fragmentation, degradation, and packaging into other particle categories. In this study, the least-processed particles were observed at the New England shelf break in late fall, where large phytodetrital aggregates dominated POC flux and contained large phytoplankton cells (primarily *Coscinodiscus*). This apparent aggregation and export of a fall diatom bloom attenuated moderately with depth (b=0.8), similar to the global average b value of 0.86 calculated by Martin et al. (1987). Phytodetrital aggregates therefore appeared to be a more efficient carbon export pathway than the long fecal pellets generated by crustaceous zooplankton.

Our observations also confirmed the episodically influential role of gelatinous zooplankton in generating sinking particles. Salp fecal pellets have long been recognized to have an outsized and episodic influence on carbon export (e.g. Iseki 1981; Bruland and Silver 1981; Caron et al. 1989), and their importance was reconfirmed during our month-long observations in the subarctic Pacific. The low numerical abundance of salp pellets increased the uncertainty of our estimates, but our observations still suggested that they attenuated little with depth, likely due to their rapid sinking speeds (>1000 m d^-1^, Phillips et al. 2009). The true effect of salp fecal pellets on carbon export is probably overlooked by many sampling programs, especially in the upper ocean where trap deployment durations are short and trap collection areas are relatively small. In the California Current, we also observed abundant large-loose pellets that were most likely generated by pyrosomes. Episodic pyrosome blooms may also play an outsized role in carbon export, similar to salps, though the influence of these pellet types has not been studied in as much detail.

These observations from distinct flux environments help to better resolve the importance of small particles (< 100 *μ*m) in the biological carbon pump. When POC fluxes were relatively high (>2 mmol C m^-2^ d^-1^), small particles consistently contributed around 5% to the total POC flux, suggesting a constant relationship with large particle flux. In low flux environments, this coupling of small particles with large particles appeared to weaken and become more variable, and their contribution to POC flux increased to 19% on average (range = 1-46%). These observations are consistent with previous estimates of carbon flux contributions by small particles in the low flux Sargasso Sea (18-78%; Durkin et al. 2015) and the higher flux Porcupine Abyssal Plain (5-25%; Bol et al. 2019).

Our observations also challenge the conventional wisdom that small particles attenuate rapidly with depth, and indicate that not all small particles are created equal. Unlike previous work (Durkin et al. 2015), we did not observe an increase in the relative contribution of all small particles to POC flux with depth. Even so, short fecal pellet fluxes either did not attenuate or increased with depth in the mesopelagic, likely due to production by mesopelagic zooplankton such as larvaceans. This observation is consistent with those of Silver and Gowing (1991). Mini pellet fluxes were elevated in the upper 150 m of the mesopelagic, and decreased at deeper depths, but are produced by microzooplankton (cilliates, dinoflagellates, phaeodarians, radiolarians) and hydromedusae (Gowing and Silver 1985) that can reside in the mesopelagic. Dense detritus did not appear to attenuate with depth, and its source is more uncertain. These particles could be fragmented or amorphous fecal pellets or just very densely packed aggregates. Some Phaeodarians could also be mistaken for dense detritus (e.g. Phaeogymnocellida, Gowing and Coale 1989; Nakamura and Suzuki 2015).

We suggest that small particles attenuated little with depth because these particles are both produced at depth and have rapid settling speeds relative to their size. McDonnell and Buesseler (2010) measured faster average sinking speeds in small particle size classes (<100 *μ*m), and attributed differences to the higher density and streamlined shapes of small particles. Similarly, Alldredge and Gotshalk (1988) measured lower sinking speeds of aggregates than would be expected from their size alone, and attributed the discrepancy to the low density and high drag of these large particles. The data presented here and in previous studies indicate that our assumptions about carbon export based on particle sizes should be reevaluated (Iversen and Lampitt 2020).

Our experience helped to identify some technical methods that we recommend for future studies. First, we found that storing gel layers in the freezer prior to imaging is a good alternative if immediate imaging at sea is not possible. In some frozen samples, gels contained air bubbles after thawing which did not always outgas. Our particle ID protocol excluded these bubbles from flux calculations, but their presence could still obscure some large particles. This problem can be avoided if the entire gel area is imaged at low magnification while at sea before freezing for additional high magnification imaging on land. Importantly, the accuracy of imaging after freezing depended on illumination method. Even in fresh samples, oblique lighting of particles was quantitatively better than brightfield illumination at detecting small and translucent particles. After freezing, particles disrupted by a freeze-thaw processes are increasingly translucent and are increasingly under-detected by brightfield illumination. One benefit of brightfield lighting is that it requires shorter image exposure times, which can make imaging in rough sea states easier. Computational processing of brightfield images is also faster because it does not require analysis of multiple color channels. However due to the improved informational content and accuracy, we recommend using oblique lighting to quantify particle POC fluxes.

Our approach to model POC fluxes from particle imagery can be improved with additional observations that pair chemical measurements of POC flux with imagery. In some locations, our model parameters did not predict variation in POC flux accurately and we do not know whether that was caused by poor model parameterization or by methodological issues during a particular cruise. For example, modeled POC flux at the New England shelf break in June did not match the measured depth attenuation profile of POC flux. A large number of amphipods were picked out of the formalin-preserved particles used to measure POC flux and may have influenced the measurement in an unknown way. Additionally, the brightfield illumination used to image the gel layers during this cruise systematically underestimated flux of translucent and small particles, which could have been an important component of the measured fluxes. For example, individual coccolithophore cells (both *Emiliania*- and *Helicosphaera*-like) and terrestrial pollen spores were qualitatively abundant within the gel layers, but could not be accurately quantified because their sizes (∼5-20 *μ*m) were below the resolution of the camera and they were relatively translucent under brightfield illumination.

More observations would also improve our parameterization of long fecal pellet carbon fluxes. Long pellets have a large influence on modeled POC fluxes but have highly variable carbon contents that are poorly constrained by a single parameterization. Long fecal pellets are produced by a variety of organisms whose fecal pellet carbon densities vary across species and with food availability (Urban-Rich et al. 1998; Atkinson et al. 2012). Our modeled POC density for these pellets (0.1 * 10^-9^ mgC *μ*m^-3^) is well within the wide range of published values but slightly higher than the value used by Wilson et al. (2008), who chose 0.08 mgC mm^-3^ as a mid-range value. Discrepancies between our modeled POC fluxes and the measured values could be caused by the large natural variation in the carbon density in these particle types when these pellets are a large fraction of the total flux, such as at the New England shelf break in June of 2016. Additional observations will help us to determine whether additional model parameters should be added and whether the existing parameters can be refined.

Current approaches that collect snapshots of carbon export in the mesopelagic cannot resolve the variability that occurs over days, weeks, and months or the episodic processes that have an outsize role in carbon export (e.g. salp fecal pellet fluxes). This study showed extreme variability in the type of particles and their contribution to flux with depth, and this variability was evident across seasons or even within a few weeks at the same location. Understanding this variability will be required to parameterize models of the biological pump. To obtain such data, we need technologies that can make sustained observations that are highly resolved in time. This study demonstrates that images of sinking particles can be used to reasonably predict the magnitude of POC flux and possibly to predict attenuation of flux with depth. This suggests that in situ imaging technologies that resolve and identify sinking particles have the potential to significantly improve quantification and monitoring of the biological carbon pump (Giering et al. 2020). Efforts to develop these instruments are already underway (Bishop et al. 2016; McGill et al. 2016; Bourne et al. 2019) and produce images analogous to those analyzed in this study. If mechanistic models of the biological pump can reasonably predict the generation of different particle classes in the euphotic zone, the type of data presented here can be used to extend those models to accurately predict carbon flux and attenuation through the mesopelagic.

## Acknowledgements

This work was supported by an NSF EAGER award to CAD (OCE-1703664), MLE (OCE-1703422), and MO (OCE-1703336), and also by the NASA EXPORTS program (80NSSC17K0662), a NASA New Investigator award to MLE (NNX14AM01G), the Rhode Island Endeavor Program (RIEP), NASA’s PACE mission, and the Schmidt Ocean Institute. We thank the captains and crews of the R/V Endeavor, R/V Falkor, and R/V Revelle who made this research possible. This work was greatly assisted and improved by the science parties of the EN572, EN581, Sea to Space, and EXPORTS expeditions, and also by Alyson Santoro, Dave Siegel, Jason Graff, Debbie Steinberg, Karen Stamieszken, Jessica Sheu, Annie Bodel, Steven Pike, Jessica Drysdale, Elly Breves, and Tom Connolly. The authors have no conflicts of interest to report.

## Supplemental Tables and Figures

**Supplemental Figure 1.**
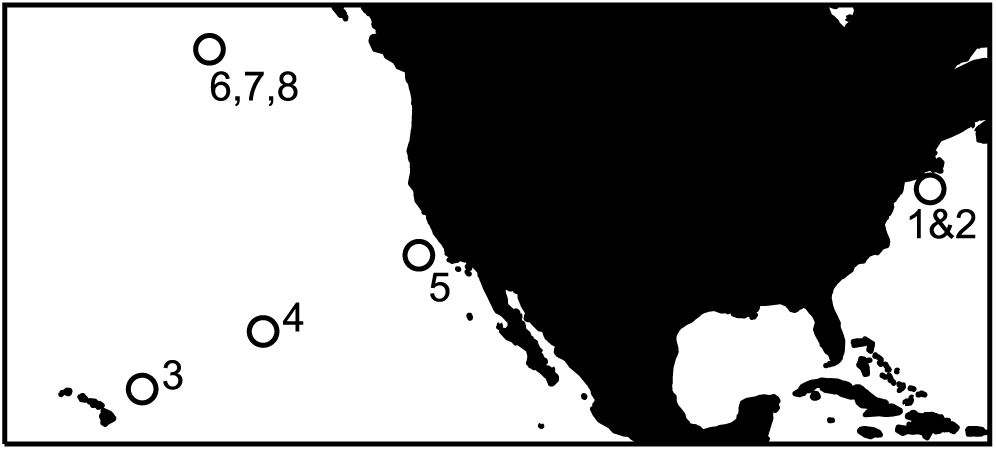
Map of sampling locations in the North Pacific Ocean (locations 3-8) and North Atlantic Ocean (locations 1-2). The continent of North America is shaded in black. Numbers correspond to locations in Table 1.

**Supplemental Figure 2.**
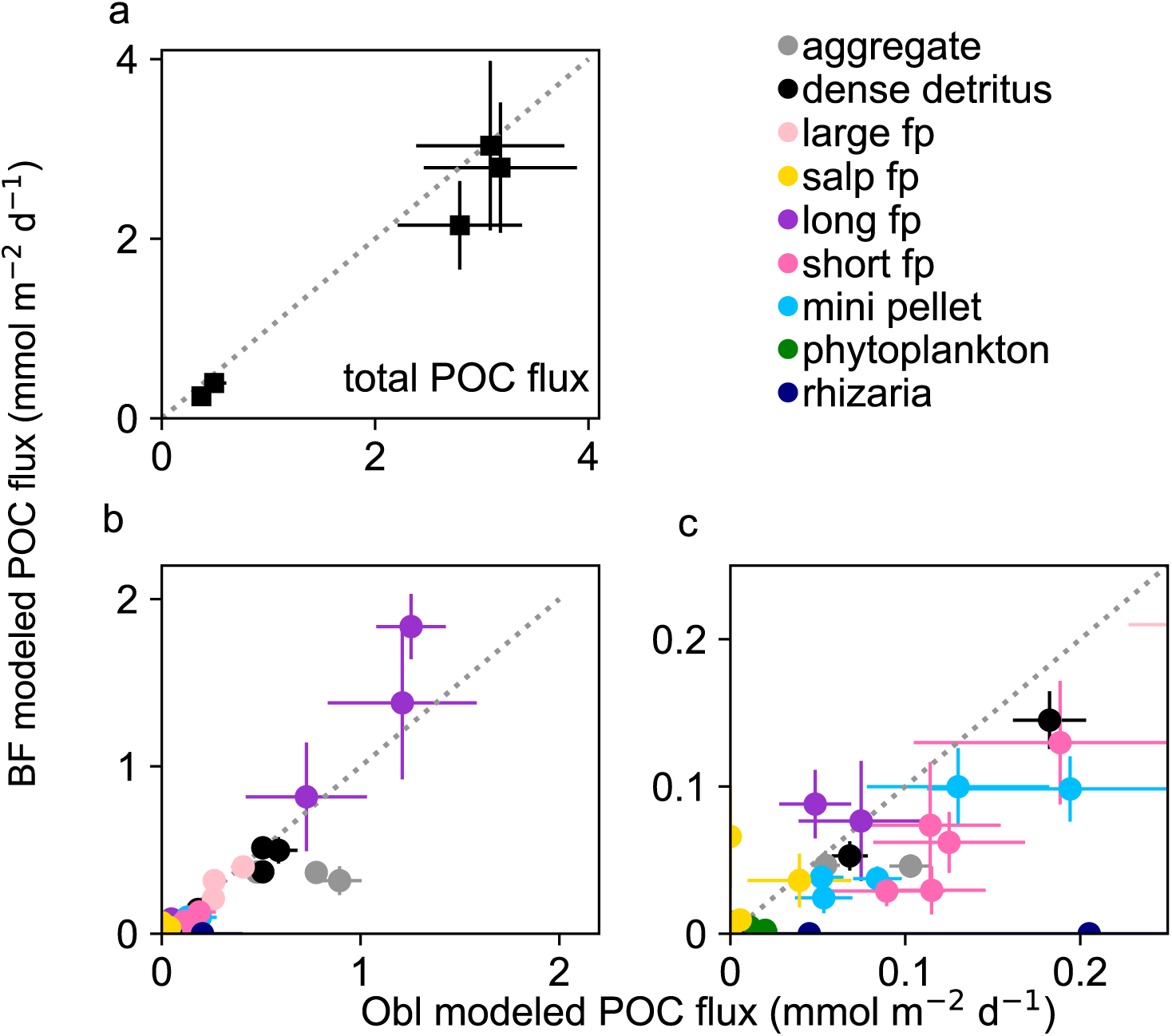
Effect of sample illumination method on detection and modeling of particulate organic carbon (POC) flux in 5 samples collected during the FK170124 cruise. a)Total modeled POC flux under brightfield (BF) versus oblique (Obl) illumination. b) Modeled POC flux of each particle type under brightfield versus oblique illumination. c) Same as panel B but scaled to highlight particles types with low POC fluxes. Error bars indicate the propagated uncertainty calculated from the number of particles counted in each classification and size bin. Dashed line indicates a 1:1 relationship. fp=fecal pellet.

**Supplemental Figure 3.**
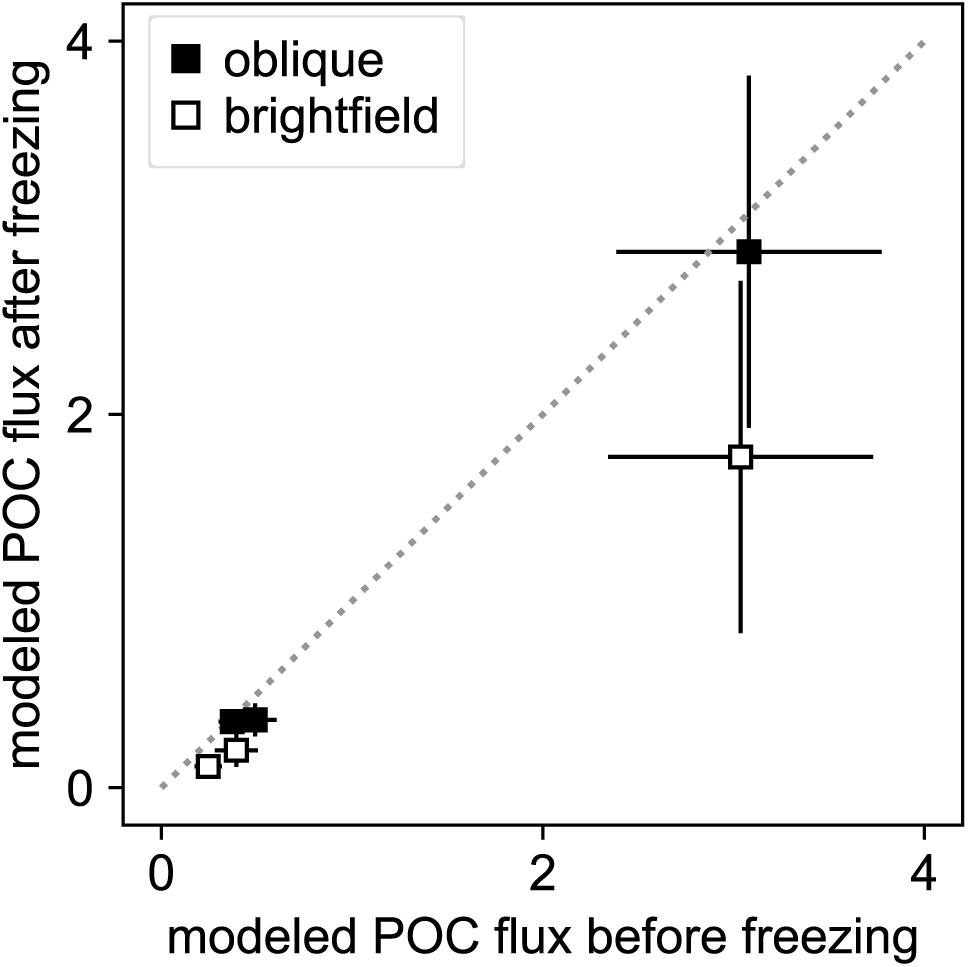
Effect of freezing gel samples on the total particulate organic carbon (POC) flux modeled from three samples collected during the FK170124 cruise. Samples were imaged immediately after collection and again after freezing and thawing under both brightfield and oblique illumination. Error bars indicate the propagated uncertainty calculated from the number of particles counted in each classification and size bin.

**Supplemental Figure 4.**
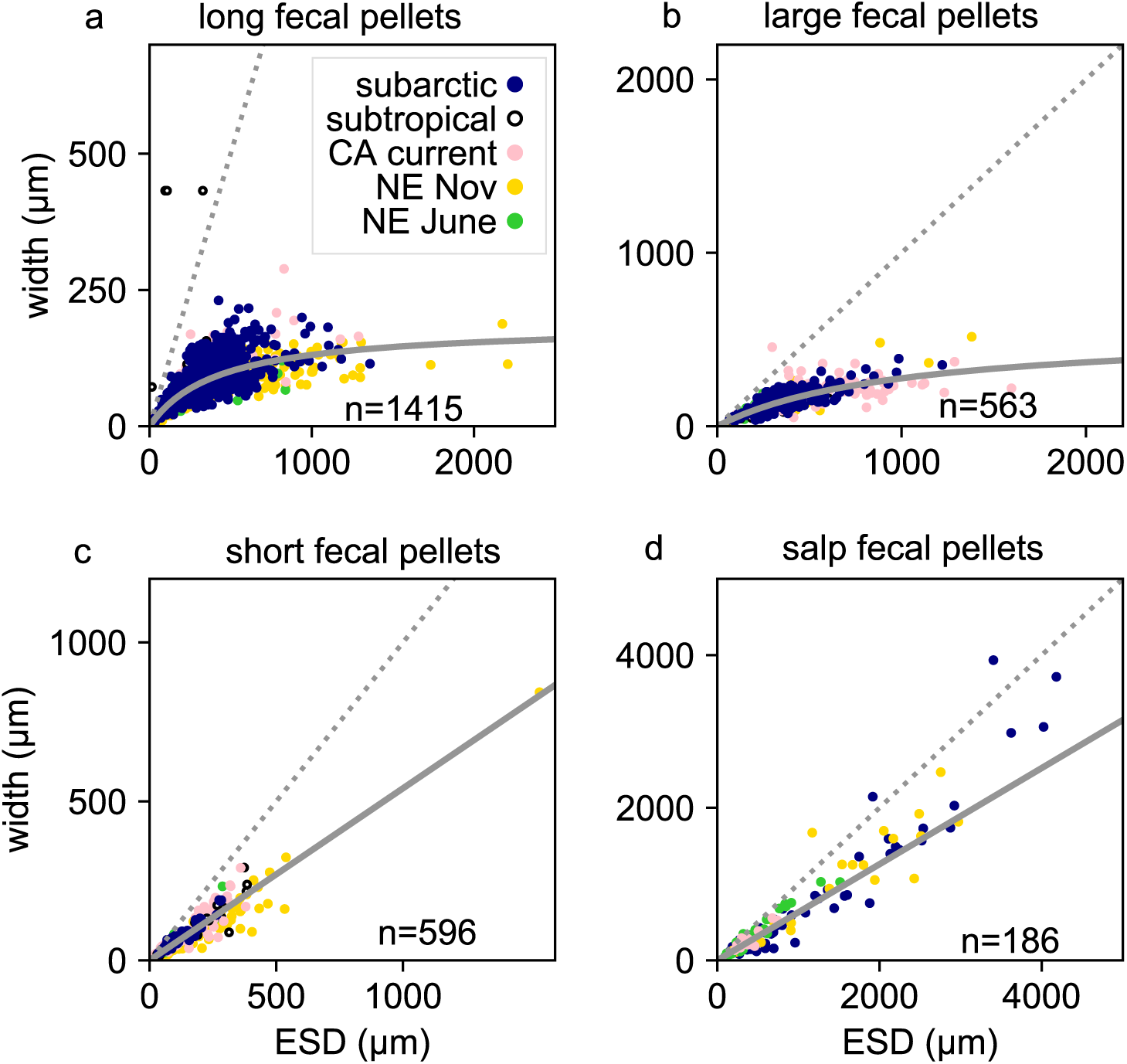
Relationship between manually measured fecal pellet widths and their equivalent spherical diameter (ESD). A representative subset (n) of a) long fecal pellets, b) large-loose fecal pellets, c) short fecal pellets, and d) salp fecal pellets were measured from all sampling locations. Dashed line represents a slope of 1, grey line is the relationship used for each pellet type. CA = California, NE= New England shelf break.

**Supplemental Figure 5.**
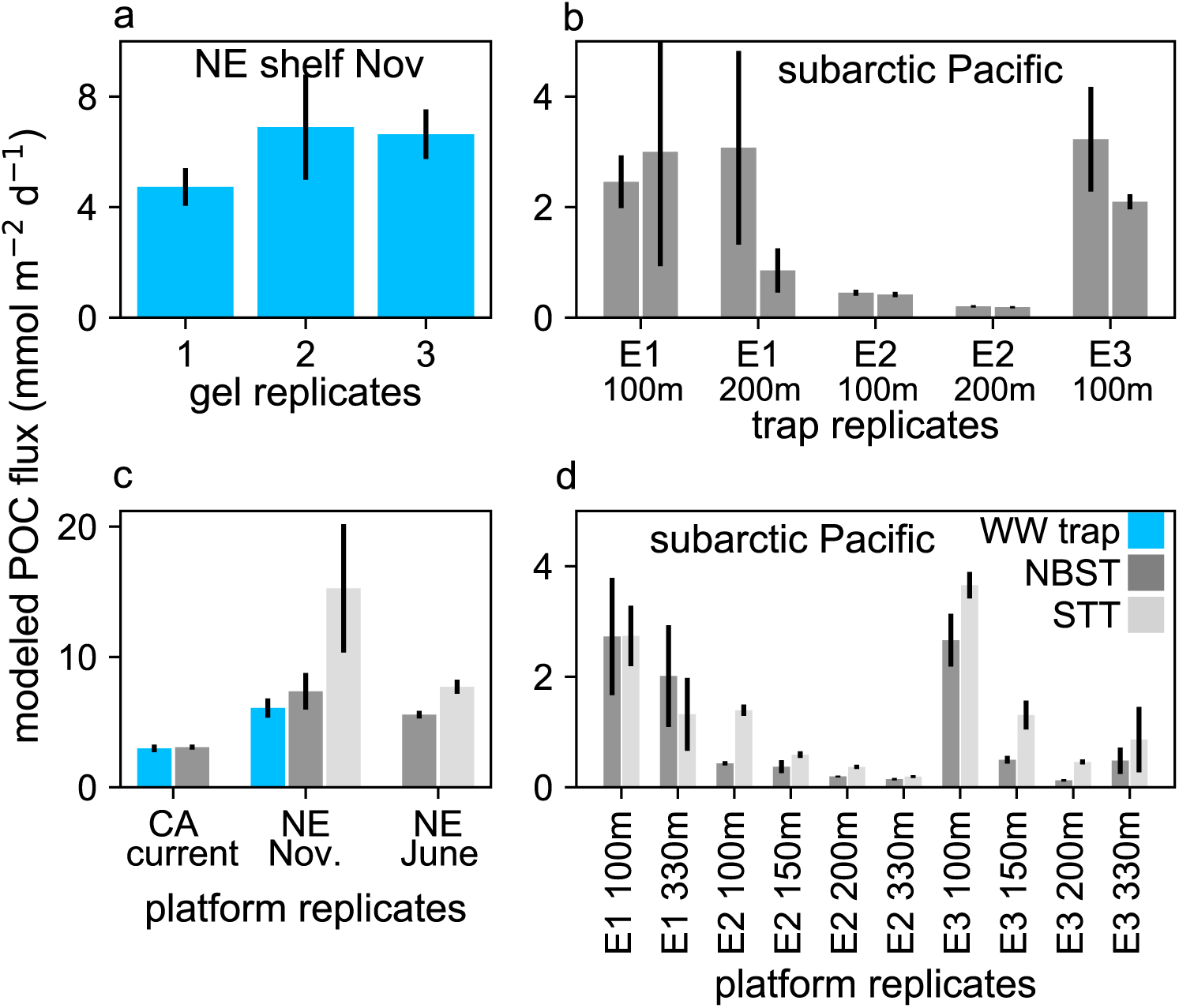
Reproducibility of modeled particulate organic carbon (POC) fluxes based on a) intraplatform stochasticity in particle collection, b) interplatform variability at small spatial scales and c&d) differences in trap platform types. a) POC flux modeled from three gel layers on the same Wirewalker (WW) trap deployed at 150 m at the New England (NE) shelf break in November. b) POC fluxes modeled from samples collected by duplicate neutrally-buoyant sediment traps (NBST) deployed at the same time and depth in the subarctic North Pacific. Deployment periods denoted by E1 (epoch 1), E2 (epoch 2), and E3 (epoch 3). c&d) POC flux modeled from samples collected by different trap platforms deployed at the same time and depth. c) Samples collected in the California (CA current) and New England shelf break. d) Samples collected in the subarctic North Pacific. Replicates of the same platform type were averaged in panels c&d. Error bars indicate the propagated uncertainty calculated from the number of particles counted in each classification and size bin.

**Supplemental Table 1.** Flux attenuation slopes (b values) of major particle types identified by the PCA. Uncertainties associated with the b value of each deployment indicate the standard deviation of randomly generated data normally distributed within individual flux uncertainties. Grey values were poorly modeled by the observations due to low observations of the particle type at the location or because their fluxes did attenuate exponentially with depth. The mean across deployments includes well constrained (bold) values and the standard deviation among those values. Location include the New England (NE) shelf break and the subarctic North Pacific (Pac) during the sampling “epochs” E1, E2, and E3.

| Deployment | aggregate | long fecal pellets | mini pellets | dense detritus | salp fecal pellets | short fecal pellets |
| --- | --- | --- | --- | --- | --- | --- |
| NE shelf Nov | **0.8±0.1** | 0.3±0.2 | **1±0.3** | **1±0.2** | no convergence | **0.9±0.4** |
| NE shelf June | 0.1±0.2 | **1.6±0.1** | 0.2±0.2 | 0.05±0.09 | 0.02±0.06 | 0.02±0.05 |
| subarctic Pac.<br>E1 | **0.5±0.05** | **1.6±0.2** | **1.1±0.2** | 0.2±0.1 | 0.1±0.2 | 0.1±0.2 |
| subarctic Pac.<br>E2 | **0.6±0.08** | **1.8±0.2** | **0.8±0.1** | 0.02±0.05 | no convergence | 0.03±0.1 |
| subarctic Pac.<br>E3 | **1.2±0.08** | **2.3±0.2** | **1.1±0.2** | 0.4±0.2 | no convergence | 0.07±0.2 |
| mean ± std | **0.8±0.3** | **1.8±0.3** | **1±0.1** | NA | NA | NA |

## Supplemental Methods

### Particulate organic carbon fluxes

Particulate organic carbon (POC) was determined using slightly different methodologies on the Endeavor, Falkor, and EXPORTS cruises because of differences in the sampling objectives of the projects. For instance, parallel collection of samples for the radionuclide ^234^Th during EXPORTS (Estapa et al., in revision) required substitution of low-background QMA filters for GF/F filters, and the availability of different analytical techniques for particulate inorganic carbon varied among projects. The reader is directed to Chaves et al. (in review) for detailed discussion of the different methods’ caveats. Particulate organic carbon collected from Endeavor deployments was determined directly by fuming dry GF/F filters over concentrated hydrochloric acid for 24 h, followed by combustion elemental analysis (Exeter Analytical CE-440). Particulate organic carbon collected from Falkor deployments was determined by difference between total carbon (TC) and particulate inorganic carbon (PIC) on different sample splits. Total carbon was determined by combustion elemental analysis of a GF/F split. Particulate inorganic carbon was determined after filtering onto polycarbonate membranes and rinsing with pH 8.5-buffered Milli-Q water. Membrane filters were dried, acidified (5% HNO_3_), and dissolved calcium and sodium were determined using flame atomic absorption spectroscopy (PerkinElmer AAnalyst 800). Particulate inorganic carbon from the equivalent calcium carbonate was computed after a small sea salt correction based on the dissolved sodium concentration. Particulate organic carbon was determined as the difference between TC and PIC. Particulate organic carbon samples collected from the EXPORTS deployments was measured similarly to the Falkor deployments as the TC-PIC difference, but PIC was determined from QMA filters after β-counting for ^234^Th fluxes, gravimetric splitting, and then coulometric analysis (Honjo et al. 2000). Bulk POC fluxes to EXPORTS traps were not used to constrain the image-based particle carbon model presented below because large numbers of swimmers were present in the EXPORTS trap deployments. Instead, the particle area fluxes to the gels were used to help perform a correction for swimmer carbon remaining in trap samples after picking, described in detail by Estapa et al. (in review). All POC flux data are available at https://seabass.gsfc.nasa.gov/archive/SKIDMORE/estapa/EXPORTS/EXPORTSNP (last accessed 6/2/2020), https://seabass.gsfc.nasa.gov/archive/SKIDMORE/estapa/Opt_Sed_Trap_Cal (last accessed 6/2/2020), and https://www.bco-dmo.org/dataset/765429 (last accessed 6/2/2020).

### Image processing and particle detection

All brightfield-illuminated images were converted to unsigned integers of 8 bits (uint8) greyscale using the skimage.color.rgb2grey function, which applies the equation Y = (0.2125×red) + (0.7154×green) +(0.0721×blue). To remove the background from each brightfield image, the median pixel value calculated from a blank gel image was subtracted from a median filtered greyscale image and this median-filtered difference was then subtracted from the uint8 greyscale image. To identify particle candidates, a threshold filter was applied to the greyscale images to select the darkest pixels, creating a binary image. All holes in the binary particle image were filled using *ndimage.morphology.binary_fill_holes* function in Python’s scipy.

Because out-of-focus particles and other noise in the image can also be detected by a simple thresholding filter of pixel brightness, additional processing steps were taken to identify only particles with in-focus outer edges. A Sobel filter was applied to the greyscale images to identify locations where large changes in pixel brightness occur over a small area (i.e. edge detection). An eroded particle mask was created from the binary image of particle candidates and was used to mask the interior pixels of the Sobel filtered image. A brightness threshold was applied to the masked, Sobel-filtered image to identify locations with sharp gradients in pixel brightness that occurred on the outer edges of particle candidates. Finally, particles were recorded if locations with in-focus edges overlapped with pixel locations of particle candidates identified in the binary image. The pixel area, location, and the Sobel-filter edge brightness of each particle were recorded (Fig. 1a).

The background of an obliquely-lit image has an intermediate pixel brightness, so particles with intermediate brightness, such as brown fecal pellets, will not be detected by a simple thresholding approach sensitive only to the brightest and darkest pixels. For this reason, color channels were also incorporated into analysis of obliquely-lit micrographs. Otherwise, the oblique analysis was identical to the processing steps used for brightfield images. To remove the background from obliquely lit images, these same background-removal calculations were applied to the greyscale, red, and blue image channels separately. To highlight brown particles with intermediate pixel brightness, the background-removed blue channel image was subtracted from the background-removed red channel image. To detect particle candidates, a threshold filter was applied to detect the brightest and darkest pixels in the greyscale image and the brightest pixels in the red channel-minus-blue-channel image. The in-focus particles were then identified and recorded in the same way as described for the brightfield images.

Some large particles were detected within multiple focal planes of the same field of view. To avoid counting the same particle more than once, additional processing steps were applied to remove duplicates from the measured particle dataset within each field of view. For each particle in an image, neighboring particles were detected within 150 pixels of the particle location’s bounding box. This neighboring area size was chosen due to the design of the microscope, whose focusing mechanism shifts the image side-to-side, and the rocking of the ship, which caused slight movement of particles while imaging. Neighboring particles were considered duplicates if their areas were within 70% of each other and they were imaged in different focal planes. The 70% area threshold was selected by manually checking a subset of the images. Even so, a small number of particles may have been counted twice while another small number of particles may have been incorrectly identified as duplicates and omitted from enumeration. The particle with the sharpest edges based on the Sobel-filtered edge brightness values was retained in the dataset for further analysis (Fig. 1b).

Measured properties of each particle included surface area (*μ*m^2^), major axis length (*μ*m), minor axis length (*μ*m), perimeter, pixel coordinates of the bounding box, and coordinates of each pixel in the particle. Equivalent spherical diameter (ESD) was also calculated from the surface area and recorded along with image file names, field of view, and focal plane. The data from every measured particle detected in a sample at a single magnification was saved as a comma-delimited text file. The particle measurements from all four magnifications were then combined and each particle image was extracted from original gel micrographs and saved into a directory of individual particle images for that sample.

