## Supplemental File 1 for "A visual tour of carbon export by sinking particles": gel_trap_image_processing_EN572_bf.py

```
#!/Users/Colleen/anaconda3/bin/python3

##Image analysis for gel trap images, brightfield lighting
##Usage: gel_trap_image_processing [input_directory] [output_mask_directory] [magnification, an integer] [blank reference image]
##scale of pixels per micron: 7x = 0.065 , 20x = 0.187 , 40x = 0.375 , 80x=0.708, 115x = 1.116

import sys, os
import numpy as np
from skimage import io, filters, color, measure, util, morphology
from scipy import ndimage as ndi
import matplotlib.pyplot as plt
from skimage.filters import sobel
import math
import pandas as pd

directory = sys.argv[1]
out_directory = sys.argv[2]
magnification = int(sys.argv[3])
ref_fh = sys.argv[4]

#set the scale of image pixels
if magnification is 7:
    scale = 0.065
    bin_index =  np.arange(14,26,1)
elif magnification is 20:
    scale= 0.187 
    bin_index =  np.arange(12,26,1)
elif magnification is 40:
    scale= 0.375  
    bin_index =  np.arange(9,26,1)
elif magnification is 80:
    scale= 0.708
    bin_index =  np.arange(6,26,1)
elif magnification is 115:
    scale= 1.116
    bin_index =  np.arange(6,26,1)
else:
    print('No scale for this magnification included in script')


#Create empty lists for every measurement property desired for a particle
count = 0
particle_area = []
particle_ESD = []
particle_perimeter = []
particle_major_axis= []
particle_minor_axis= []
particle_numberID = []
file_name = []
field = []
fplane = []
brightest_edge = []
particle_coordinates = []
bounding_box = []

#Read the blank reference photo, convert to grey scale, calculate the median intensity value of grey scale, and the red and blue channels.
photo_blank = io.imread(ref_fh, plugin= 'pil')
photo_blank_grey1 = color.rgb2grey(photo_blank) #converts to luminescence units, scale -1 to 1 (Y = 0.2125 R + 0.7154 G + 0.0721 B)
photo_blank_grey=util.img_as_ubyte(photo_blank_grey1)  #converts to a 8-bit ubyte, scale 0 to 255

median_grey_value=np.median(photo_blank_grey)

##Read image, convert to greyscale, and remove the background by subtracting the regional median (calculated for each pixel as the median of the surrounding pixels) and normalizing to the blank image median value##
#Now have background-removed image
for file in os.listdir(directory):
    photo_path = os.path.join(directory , file)
    try:
        photo = io.imread(photo_path, plugin='pil')
    except:
        continue
    photo_grey1 = color.rgb2grey(photo) #converts to luminescence units, scale -1 to 1 (Y = 0.2125 R + 0.7154 G + 0.0721 B)
    photo_grey=util.img_as_ubyte(photo_grey1) #converts to a 8-bit ubyte, scale 0 to 255
    if magnification is 7 or magnification is 20:
    	photo_grey_median = filters.median(photo_grey,selem=morphology.disk(200)) #disk shape over which median value is calculated must be larger than the largest particle
    if magnification is 40 or magnification is 80 or magnification is 115:
    	photo_grey_median = filters.median(photo_grey,selem=morphology.disk(500)) #disk shape over which median value is calculated must be larger than the largest particle
    photo_grey_median_diff = photo_grey_median - median_grey_value #difference between median filtered image and the median value of a blank image.  Use this to indicate variation in the background intensity across the image space.
    photo_grey_nobg = photo_grey-photo_grey_median_diff #background subtracted from the grey-scale image.  This step is especially necessary if the lighting is not even across the entire image (e.g. top of image is brighter than the bottom of the image)

    ##Define the marker pixels of all possible particles##
    #need to change the theshold value depending on magnification because lighting gets dimmer as magnification gets higher
    markers = np.zeros_like(photo_grey_nobg)
    if magnification is 7:
    	markers[photo_grey_nobg <= 150] = 1
    	markers[photo_grey_nobg > 150] = 0
    if magnification is 20:
    	markers[photo_grey_nobg <= 165] = 1
    	markers[photo_grey_nobg > 165] = 0
    if magnification is 40:
    	markers[photo_grey_nobg <= 165] = 1
    	markers[photo_grey_nobg > 165] = 0
    if magnification is 80:
    	markers[photo_grey_nobg <= 155] = 1
    	markers[photo_grey_nobg > 155] = 0
    if magnification is 115:
        markers[photo_grey_nobg <= 185] = 1
        markers[photo_grey_nobg > 185] = 0
    
    markers_fill = ndi.morphology.binary_fill_holes(markers)
    labeled_particles, _ = ndi.label(markers_fill)

    #Create a mask of insides of the particles, identify edges from sobel filter and mask out the inside of particles
    #Need to mask out center of particles so edges are not detected on the interior of the particle (only want to detect particles whose outside edges are sharp).
    particle_erode = morphology.erosion(markers_fill, selem = morphology.disk(1))
    elevation_map = sobel(photo_grey_nobg)
    elevation_map[particle_erode==True]=0

    ##Identify sharpest edges from the sobel filter##
    #Also need to change the threshold of what is concidered "sharp" as magnification increases because more difficult to get crisp images as you zoom in(especially on a rocking ship)
    edges = np.zeros_like(photo_grey_nobg)
    if magnification is 7:
    	edges[elevation_map <= 10] = 0
    	edges[elevation_map > 10] = 255
    if magnification is 20:
    	edges[elevation_map <= 10] = 0
    	edges[elevation_map > 10] = 255
    if magnification is 40:
    	edges[elevation_map <= 10] = 0
    	edges[elevation_map > 10] = 255
    if magnification is 80:
    	edges[elevation_map <= 10] = 0
    	edges[elevation_map > 10] = 255
    if magnification is 115:
    	edges[elevation_map <= 6] = 0
    	edges[elevation_map > 6] = 255
    labeled_edges , _ = ndi.label(edges)

    ##Identify only in-focus particles by particle indexes that overlap with edge indexes##
    infocus_object_img = np.zeros_like(labeled_particles)
    edge_properties = measure.regionprops(labeled_edges)
    particle_properties = measure.regionprops(labeled_particles)
    infocus_index=[]
    for edge in edge_properties:
        e_coords = edge.coords
        for x in e_coords:
            infocus = labeled_particles[x[0],x[1]]
            if infocus not in infocus_index:
                infocus_index.append(infocus)
            
    for y in infocus_index:
        infocus_object_img[labeled_particles==y] = y

    #Measure and record each particle identified
    scale_area = scale**2   #square pixels per square micron
    properties = measure.regionprops(infocus_object_img)
    for x in properties:
        count = count +1
        px_area=x.area
        um_area = px_area / scale_area
        min_axis = x.minor_axis_length / scale
        maj_axis = x.major_axis_length / scale
        perim = x.perimeter / scale
        ESD = 2*(math.sqrt(um_area/math.pi))
        particle_perimeter.append(perim)
        particle_major_axis.append(maj_axis)
        particle_minor_axis.append(min_axis)
        particle_area.append(um_area)
        particle_ESD.append(ESD)
        particle_numberID.append(count)
        file_name.append(file)
        name_words = file.split('_')
        focalplane = name_words[len(name_words)-1]
        if focalplane[0:2].isdigit():
        	fov = focalplane[0:2]
        	plane = focalplane[2]
        else:
        	fov = focalplane[0]
        	plane = focalplane[1]
        field.append(fov)
        fplane.append(plane)
        edge_area=elevation_map[x.bbox[0]:x.bbox[2],x.bbox[1]:x.bbox[3]]
        max_edge_intensity = np.max(edge_area)
        brightest_edge.append(max_edge_intensity)
        particle_coordinates.append(np.ndarray.tolist(x.coords))
        bounding_box.append(str([x.bbox[0],x.bbox[2],x.bbox[1],x.bbox[3]]))

    print('File ' + file + ' has been processed.')

#Input all data into a dataframe.#
data = pd.DataFrame(np.stack((particle_numberID, particle_area, particle_ESD, particle_minor_axis, particle_major_axis, particle_perimeter, file_name, field, fplane, brightest_edge, particle_coordinates, bounding_box),-1),columns=['Number','Area','ESD','minor_length','major_length','perimeter','file_name', 'fov','focal_plane','edge_intensity','particle_coordinates','bounding_box'])
data.to_csv(out_directory + 'All_particles_' + str(magnification) + 'x.csv')

print('Images have been processed.  Now deduplicating particle data to save.')

#make a directory to save all the individual particle images into
#os.mkdir(os.path.join(out_directory,'Detected_particles'))

Deduplicated_data = pd.DataFrame()


for field in data.fov.unique():
    field_data = data[data.fov == field]
    bbox_coordinates_list = pd.DataFrame(columns=['Top','Bottom','Left','Right'])
    for a in field_data.index:
        bbox_coordinates = field_data.loc[a].bounding_box[1:(len(field_data.loc[a].bounding_box)-1)].split(', ')
        bbox_coordinates_list.loc[a]=[int(bbox_coordinates[0]),int(bbox_coordinates[1]),int(bbox_coordinates[2]),int(bbox_coordinates[3])]
    field_data = pd.concat([field_data,bbox_coordinates_list], axis = 1)
    for b in field_data.Number:
        b_row = field_data.loc[field_data.Number == b]
        b_row.Area = b_row.Area.astype('float64')
        topside=b_row.Top.values[0]-150
        bottomside=b_row.Bottom.values[0]+150
        leftside=b_row.Left.values[0]-150
        rightside=b_row.Right.values[0]+150
        if topside <= 0:
            topside=0
        if bottomside >= np.shape(photo)[0]:
            bottomside = np.shape(photo)[0]
        if leftside <= 0:
            leftside = 0
        if rightside >= np.shape(photo)[1]:
            rightside = np.shape(photo)[1]
        surrounding_box=[int(topside),int(bottomside),int(leftside),int(rightside)]
        current_fp = b_row.focal_plane
        particles_in_other_fp = field_data[field_data.focal_plane != current_fp.values[0]]
        overlap_1 = particles_in_other_fp[particles_in_other_fp.Top >= surrounding_box[0]]
        overlap_2 = overlap_1[overlap_1.Bottom <= surrounding_box[1]]
        overlap_3 = overlap_2[overlap_2.Left >= surrounding_box[2]]
        potential_duplicates = overlap_3[overlap_3.Right <= surrounding_box[3]]
        potential_duplicates1 = potential_duplicates[potential_duplicates.Area <= 1.42*b_row.Area.values[0]]
        potential_duplicates2 = potential_duplicates1[potential_duplicates1.Area >= 0.7*b_row.Area.values[0]]       
        duplicates = pd.DataFrame(columns = potential_duplicates2.columns)
        duplicates = duplicates.append(b_row)
        for x in np.unique(potential_duplicates2.focal_plane):
            fp_x = potential_duplicates2[potential_duplicates2.focal_plane == x]
            fp_x.Area = fp_x.Area.astype('float64')
            if len(fp_x.Number) > 1:                                                   #find which ones are in the same focal plane
                Area_difference = abs(fp_x.Area - (b_row.Area.values[0]))              #identify which one is closest in size to the b_row particle
                Area_difference = Area_difference.astype('float64')
                selected_duplicate_index = Area_difference.idxmin(axis = 0)
                duplicates = duplicates.append(potential_duplicates2.loc[selected_duplicate_index])   #add the true duplicate from that focal plane to the list of duplicates
            else:                                      #for all other duplicates, that were the only one identified in their focal plane:
                duplicates = duplicates.append(fp_x)   #add the duplicate from that focal plane to the list of duplicates
        duplicates.edge_intensity = duplicates.edge_intensity.astype('float64')
        in_focus_particle = duplicates.edge_intensity.idxmax(axis = 0)
        for p in duplicates.index:
            if p in Deduplicated_data.index:
                Deduplicated_data=Deduplicated_data.drop(p) 
        Deduplicated_data=Deduplicated_data.append(field_data.loc[in_focus_particle])

Deduplicated_data.to_csv(out_directory + 'Measured_particles_' + str(magnification) + 'x.csv')
Deduplicated_data = pd.DataFrame.from_csv(out_directory + 'Measured_particles_' + str(magnification) + 'x.csv')

for field in Deduplicated_data.fov.unique():
    field_data = Deduplicated_data[Deduplicated_data.fov == field]
    dedup_particles_mask = np.zeros(np.shape(photo[:,:,1]),dtype=int)
    for a in field_data.index:
        xrow = Deduplicated_data.loc[a]
        coordinates=xrow.particle_coordinates[1:len(xrow.particle_coordinates)-1]
        coordinates_list = coordinates[1:len(coordinates)-1].split('], [')
        for coordinate in coordinates_list:
            x_y_coordinates = coordinate.split(', ')
            dedup_particles_mask[int(x_y_coordinates[0]),int(x_y_coordinates[1])]=255
    io.imsave(os.path.join(out_directory , 'Measured_Particles_' + str(magnification) + 'x_Mask' + str(int(field)) + '.jpg'), dedup_particles_mask,plugin= 'pil')
    
```
