## Supplemental File 1 for "A visual tour of carbon export by sinking particles": gel_trap_image_processing_EXPORTS.py

```
#!/Users/colleen/anaconda3/bin/python

##Image analysis for gel trap images, oblique lighting
##Usage: gel_trap_image_processing [input_directory] [output_mask_directory] [magnification, an integer] [blank reference image]
##scale of pixels per micron: 7x = 0.1268 , 20x = 0.364 , 50x = 0.928 , 115x = 2.179

import sys, os
import numpy as np
from skimage import io, filters, color, measure, util, morphology
from scipy import ndimage as ndi
import matplotlib.pyplot as plt
from skimage.filters import sobel
import math
import pandas as pd

directory = sys.argv[1]
out_directory = sys.argv[2]
magnification = int(sys.argv[3])
ref_fh = sys.argv[4]

#set the scale of image pixels per micron
if magnification is 7:
    scale = 0.1268
    bin_index =  np.arange(14,26,1)
elif magnification is 20:
    scale= 0.364 
    bin_index =  np.arange(12,26,1)
elif magnification is 50:
    scale= 0.928  
    bin_index =  np.arange(9,26,1)
elif magnification is 115:
    scale= 2.179
    bin_index =  np.arange(6,26,1)
else:
    print('No scale for this magnification included in script')


#Create empty lists for every measurement property desired for a particle
count = 0
particle_area = []
particle_ESD = []
particle_perimeter = []
particle_major_axis= []
particle_minor_axis= []
particle_numberID = []
file_name = []
field = []
fplane = []
brightest_edge = []
particle_coordinates = []
bounding_box = []

#Read the blank reference photo, convert to grey scale, calculate the median intensity value of grey scale, and the red and blue channels.
photo_blank = io.imread(ref_fh, plugin= 'pil')
photo_blank_grey1 = color.rgb2grey(photo_blank) #converts to luminescence units, scale -1 to 1 (Y = 0.2125 R + 0.7154 G + 0.0721 B)
photo_blank_grey=util.img_as_ubyte(photo_blank_grey1)  #converts to a 8-bit ubyte, scale 0 to 255
photo_blank_red1 = photo_blank[:,:,0]
photo_blank_red=util.img_as_ubyte(photo_blank_red1)
photo_blank_blue1 = photo_blank[:,:,2]
photo_blank_blue=util.img_as_ubyte(photo_blank_blue1)

median_grey_value=np.median(photo_blank_grey)
median_red_value=np.median(photo_blank_red)
median_blue_value=np.median(photo_blank_blue)

##Read image, convert to greyscale, and remove the background by subtracting the regional median (calculated for each pixel as the median of the surrounding pixels) and normalizing to the blank image median value##
##Additionally, create a proxy image that highlights light brown color.  Subtracting the blue channel image from the red channel makes light brown color pixels relatively bright.
##To create light-brown proxy image, remove background in red and blue image channels in the same way as was performed for the grey scale image.  Subtract the red and blue channels (background removed) from one another.
#Now have two background-removed images: greyscale image and the light-brown proxy image.
for file in os.listdir(directory):
    photo_path = os.path.join(directory , file)
    try:
        photo = io.imread(photo_path, plugin='pil')
    except:
        continue
    photo_grey1 = color.rgb2grey(photo) #converts to luminescence units, scale -1 to 1 (Y = 0.2125 R + 0.7154 G + 0.0721 B)
    photo_grey=util.img_as_ubyte(photo_grey1) #converts to a 8-bit ubyte, scale 0 to 255
    photo_red1 = photo[:,:,0]
    photo_red=util.img_as_ubyte(photo_red1)
    photo_blue1 = photo[:,:,2]
    photo_blue=util.img_as_ubyte(photo_blue1)
    if magnification is 7 or magnification is 20:
    	photo_grey_median = filters.median(photo_grey,selem=morphology.disk(200)) #disk shape over which median value is calculated must be larger than the largest particle
    	photo_red_median = filters.median(photo_red,selem=morphology.disk(200))
    	photo_blue_median = filters.median(photo_blue,selem=morphology.disk(200))
    if magnification is 50 or magnification is 115:
    	photo_grey_median = filters.median(photo_grey,selem=morphology.disk(500)) #disk shape over which median value is calculated must be larger than the largest particle
    	photo_red_median = filters.median(photo_red,selem=morphology.disk(500)) 
    	photo_blue_median = filters.median(photo_blue,selem=morphology.disk(500))
    photo_grey_median_diff = photo_grey_median - median_grey_value #difference between median filtered image and the median value of a blank image.  Use this to indicate variation in the background intensity across the image space.
    photo_grey_nobg = photo_grey-photo_grey_median_diff #background subtracted from the grey-scale image.  This step is especially necessary if the lighting is not even across the entire image (e.g. top of image is brighter than the bottom of the image)
    photo_red_median_diff = photo_red_median - median_red_value
    photo_blue_median_diff = photo_blue_median - median_blue_value
    photo_red_nobg = photo_red-photo_red_median_diff
    photo_blue_nobg = photo_blue-photo_blue_median_diff
    photo_brown_nobg= photo_red_nobg - photo_blue_nobg #red minus blue channel (background removed) creates an image that highlights light brown pixels.

    ##Define the marker pixels of all possible particles##
    #need to change the theshold value depending on magnification because lighting gets dimmer as magnification gets higher
    #In greyscale image, brightest pixels capture white color and darkest pixels capture the dark brown/black particles.  Medium intensity colors like light brown are not distinguishable from the background.
    #In brown proxy image, brightest pixels correspond with light brown parts of particles.
    markers = np.zeros_like(photo_grey_nobg)
    if magnification is 7:
    	markers[photo_grey_nobg >= 58] = 1
    	markers[photo_grey_nobg <= 30] = 1
    	markers[photo_brown_nobg > 5] = 1
    if magnification is 20:
    	markers[photo_grey_nobg >= 58] = 1
    	markers[photo_grey_nobg <= 30] = 1
    	markers[photo_brown_nobg > 10] = 1
    if magnification is 50:
    	markers[photo_grey_nobg >= 56] = 1
    	markers[photo_grey_nobg <= 33] = 1
    	markers[photo_brown_nobg >= 9] = 1
    if magnification is 115:
        markers[photo_grey_nobg >= 60] = 1
        markers[photo_grey_nobg <= 40] = 1
        markers[photo_brown_nobg >=18] = 1
    
    markers_fill = ndi.morphology.binary_fill_holes(markers)
    labeled_particles, _ = ndi.label(markers_fill)

    #Create a mask of insides of the particles, identify edges from sobel filter and mask out the inside of particles
    #Need to mask out center of particles so edges are not detected on the interior of the particle (only want to detect particles whose outside edges are sharp).
    ##This masking step also helps when dealing with oblique images, where color is more apparent and can make "edges" appear when colors change but image is still blurry.
    particle_erode = morphology.erosion(markers_fill, selem = morphology.disk(1))
    elevation_map = sobel(photo_grey_nobg)
    elevation_map[particle_erode==True]=0

    ##Identify sharpest edges from the sobel filter##
    #Also need to change the threshold of what is concidered "sharp" as magnification increases because more difficult to get crisp images as you zoom in(especially on a rocking ship)
    edges = np.zeros_like(photo_grey_nobg)
    if magnification is 7 or magnification is 20:
    	edges[elevation_map > 8] = 255
    if magnification is 50:
    	edges[elevation_map > 8] = 255
    if magnification is 115:
    	edges[elevation_map > 9] = 255
    labeled_edges , _ = ndi.label(edges)

    ##Identify only in-focus particles by particle indexes that overlap with edge indexes##
    infocus_object_img = np.zeros_like(labeled_particles)
    edge_properties = measure.regionprops(labeled_edges)
    particle_properties = measure.regionprops(labeled_particles)
    infocus_index=[]
    for edge in edge_properties:
        e_coords = edge.coords
        for x in e_coords:
            infocus = labeled_particles[x[0],x[1]]
            if infocus not in infocus_index:
                infocus_index.append(infocus)
            
    for y in infocus_index:
        infocus_object_img[labeled_particles==y] = y

    #Measure and record each particle identified
    scale_area = scale**2   #square pixels per square micron
    properties = measure.regionprops(infocus_object_img)
    for x in properties:
        count = count +1
        px_area=x.area
        um_area = px_area / scale_area
        min_axis = x.minor_axis_length / scale
        maj_axis = x.major_axis_length / scale
        perim = x.perimeter / scale
        ESD = 2*(math.sqrt(um_area/math.pi))
        particle_perimeter.append(perim)
        particle_major_axis.append(maj_axis)
        particle_minor_axis.append(min_axis)
        particle_area.append(um_area)
        particle_ESD.append(ESD)
        particle_numberID.append(count)
        file_name.append(file)
        name_words = file.split('_')
        focalplane = name_words[len(name_words)-1]
        if focalplane[0:2].isdigit():
        	fov = focalplane[0:2]
        	plane = focalplane[2]
        else:
        	fov = focalplane[0]
        	plane = focalplane[1]
        field.append(fov)
        fplane.append(plane)
        edge_area=elevation_map[x.bbox[0]:x.bbox[2],x.bbox[1]:x.bbox[3]]
        max_edge_intensity = np.max(edge_area)
        brightest_edge.append(max_edge_intensity)
        particle_coordinates.append(np.ndarray.tolist(x.coords))
        bounding_box.append(str([x.bbox[0],x.bbox[2],x.bbox[1],x.bbox[3]]))

    print('File ' + file + ' has been processed.')

#Input all data into a dataframe.#
data = pd.DataFrame(np.stack((particle_numberID, particle_area, particle_ESD, particle_minor_axis, particle_major_axis, particle_perimeter, file_name, field, fplane, brightest_edge, particle_coordinates, bounding_box),-1),columns=['Number','Area','ESD','minor_length','major_length','perimeter','file_name', 'fov','focal_plane','edge_intensity','particle_coordinates','bounding_box'])
data.to_csv(out_directory + 'All_particles_' + str(magnification) + 'x.csv')

print('Images have been processed.  Now deduplicating particle data to save.')

#make a directory to save all the individual particle images into
#os.mkdir(os.path.join(out_directory,'Detected_particles'))

Deduplicated_data = pd.DataFrame()


for field in data.fov.unique():
    field_data = data[data.fov == field]
    bbox_coordinates_list = pd.DataFrame(columns=['Top','Bottom','Left','Right'])
    for a in field_data.index:
        bbox_coordinates = field_data.loc[a].bounding_box[1:(len(field_data.loc[a].bounding_box)-1)].split(', ')
        bbox_coordinates_list.loc[a]=[int(bbox_coordinates[0]),int(bbox_coordinates[1]),int(bbox_coordinates[2]),int(bbox_coordinates[3])]
    field_data = pd.concat([field_data,bbox_coordinates_list], axis = 1)
    for b in field_data.Number:
        b_row = field_data.loc[field_data.Number == b]
        topside=b_row.Top.values[0]-150
        bottomside=b_row.Bottom.values[0]+150
        leftside=b_row.Left.values[0]-150
        rightside=b_row.Right.values[0]+150
        if topside <= 0:
            topside=0
        if bottomside >= np.shape(photo)[0]:
            bottomside = np.shape(photo)[0]
        if leftside <= 0:
            leftside = 0
        if rightside >= np.shape(photo)[1]:
            rightside = np.shape(photo)[1]
        surrounding_box=[int(topside),int(bottomside),int(leftside),int(rightside)]
        current_fp = b_row.focal_plane
        particles_in_other_fp = field_data[field_data.focal_plane != current_fp.values[0]]
        overlap_1 = particles_in_other_fp[particles_in_other_fp.Top >= surrounding_box[0]]
        overlap_2 = overlap_1[overlap_1.Bottom <= surrounding_box[1]]
        overlap_3 = overlap_2[overlap_2.Left >= surrounding_box[2]]
        potential_duplicates = overlap_3[overlap_3.Right <= surrounding_box[3]]
        potential_duplicates1 = potential_duplicates[potential_duplicates.Area <= 1.42*b_row.Area.values[0]]
        potential_duplicates2 = potential_duplicates1[potential_duplicates1.Area >= 0.7*b_row.Area.values[0]]       
        duplicates = pd.DataFrame(columns = potential_duplicates2.columns)
        duplicates = duplicates.append(b_row)
        for x in np.unique(potential_duplicates2.focal_plane):
            fp_x = potential_duplicates2[potential_duplicates2.focal_plane == x]
            if len(fp_x.Number) > 1:                                                   #find which ones are in the same focal plane
                Area_difference = abs(fp_x.Area - (b_row.Area.values[0]))              #identify which one is closest in size to the b_row particle
                selected_duplicate_index = np.argmin(Area_difference)
                duplicates = duplicates.append(potential_duplicates2.loc[selected_duplicate_index])   #add the true duplicate from that focal plane to the list of duplicates
            else:                                      #for all other duplicates, that were the only one identified in their focal plane:
                duplicates = duplicates.append(fp_x)   #add the duplicate from that focal plane to the list of duplicates
        in_focus_particle = np.argmax(duplicates.edge_intensity)
        for p in duplicates.index:
            if p in Deduplicated_data.index:
                Deduplicated_data=Deduplicated_data.drop(p) 
        Deduplicated_data=Deduplicated_data.append(field_data.loc[in_focus_particle])

Deduplicated_data.to_csv(out_directory + 'Measured_particles_' + str(magnification) + 'x.csv')
Deduplicated_data = pd.DataFrame.from_csv(out_directory + 'Measured_particles_' + str(magnification) + 'x.csv')

for field in Deduplicated_data.fov.unique():
    field_data = Deduplicated_data[Deduplicated_data.fov == field]
    dedup_particles_mask = np.zeros(np.shape(photo[:,:,1]),dtype=int)
    for a in field_data.index:
        xrow = Deduplicated_data.loc[a]
        coordinates=xrow.particle_coordinates[1:len(xrow.particle_coordinates)-1]
        coordinates_list = coordinates[1:len(coordinates)-1].split('], [')
        for coordinate in coordinates_list:
            x_y_coordinates = coordinate.split(', ')
            dedup_particles_mask[int(x_y_coordinates[0]),int(x_y_coordinates[1])]=255
    io.imsave(os.path.join(out_directory , 'Measured_Particles_' + str(magnification) + 'x_Mask' + str(int(field)) + '.tiff'), dedup_particles_mask,plugin= 'pil')
    
```
