## Supplemental File 1 for "A visual tour of carbon export by sinking particles": Extract_particle_images_and_combine_csv_EN572.py

```
#!/Users/colleen/anaconda3/bin/python

#Endeavor EN72 brightfield lighting samples
##Extract individual particle images detected by image processing in all magnifications and recorded in one combined csv file

#Use: Extract_particles_images_and_combine_csv.py [Sample name] [image processing directory] [gel image directory]

#the image processing directory contains subfolders for each magnification that is "./Sample/Sample_7x_mask/Measured_particles_7x.csv"
#where Sample is the name of the sample and a subfolder exists for each magnification.

#The gel image directory contains subfolder for each sample: "./Sample/Sample_7x_images/" for each magnification where "Sample"=sample name

#The smallest size "particles" detected in any magnification are not identifiable because they can not be resolved by the camera
#Only extract and try to identify particles that stand a chance of being identified, otherwise will be overwhelmed by noise outside optimal size ranges
#7x image size range: >= 256 um ESD
#20x image size range: >=64 um ESD
#40x image size range: >=22 um ESD
#115x image size range: >=10 um ESD

import sys, os
import numpy as np
from skimage import io, color, measure, util, morphology
from scipy import ndimage as ndi
import math
import pandas as pd

Sample = sys.argv[1]
directory = sys.argv[2]
image_directory = sys.argv[3]

directory_7x = Sample + '_7x_mask'
directory_20x = Sample + '_20x_mask'
directory_40x = Sample + '_40x_mask'
directory_115x = Sample + '_115x_mask'
os.mkdir(os.path.join(directory,Sample,Sample + '_particle_images'))

file_7x = pd.read_csv(os.path.join(directory,Sample,directory_7x,'Measured_particles_7x.csv'))
file_20x = pd.read_csv(os.path.join(directory,Sample,directory_20x,'Measured_particles_20x.csv'))
file_40x = pd.read_csv(os.path.join(directory,Sample,directory_40x,'Measured_particles_40x.csv'))
file_115x = pd.read_csv(os.path.join(directory,Sample,directory_115x,'Measured_particles_115x.csv'))

subset_7x = file_7x.loc[file_7x.ESD >=256]
subset_20x = file_20x.loc[file_20x.ESD >=64]
subset_40x = file_40x.loc[file_40x.ESD >=45]
subset_115x = file_115x.loc[file_115x.ESD >=11]


for x in np.unique(subset_7x.file_name):
	photo = io.imread(os.path.join(image_directory,Sample,Sample+'_7x_images',x), plugin= 'pil')
	photo_subset_7x = subset_7x.loc[subset_7x.file_name == x]
	for p in photo_subset_7x.index:
		particle = photo[int(photo_subset_7x.loc[p].Top):int(photo_subset_7x.loc[p].Bottom),int(photo_subset_7x.loc[p].Left):int(photo_subset_7x.loc[p].Right)]
		particle_name1 = x.split('.jpg')
		particle_name = particle_name1[0] + '_' + str(int(photo_subset_7x.loc[p].Number)) + '.jpg'
		io.imsave(os.path.join(directory,Sample,Sample + '_particle_images',particle_name),particle)
		
for x in np.unique(subset_20x.file_name):
	photo = io.imread(os.path.join(image_directory,Sample,Sample+'_20x_images',x), plugin= 'pil')
	photo_subset_20x = subset_20x.loc[subset_20x.file_name == x]
	for p in photo_subset_20x.index:
		particle = photo[int(photo_subset_20x.loc[p].Top):int(photo_subset_20x.loc[p].Bottom),int(photo_subset_20x.loc[p].Left):int(photo_subset_20x.loc[p].Right)]
		particle_name1 = x.split('.jpg')
		particle_name = particle_name1[0] + '_' + str(int(photo_subset_20x.loc[p].Number)) + '.jpg'
		io.imsave(os.path.join(directory,Sample,Sample + '_particle_images',particle_name),particle)

for x in np.unique(subset_40x.file_name):
	photo = io.imread(os.path.join(image_directory,Sample,Sample+'_40x_images',x), plugin= 'pil')
	photo_subset_40x = subset_40x.loc[subset_40x.file_name == x]
	for p in photo_subset_40x.index:
		particle = photo[int(photo_subset_40x.loc[p].Top):int(photo_subset_40x.loc[p].Bottom),int(photo_subset_40x.loc[p].Left):int(photo_subset_40x.loc[p].Right)]
		particle_name1 = x.split('.jpg')
		particle_name = particle_name1[0] + '_' + str(int(photo_subset_40x.loc[p].Number)) + '.jpg'
		io.imsave(os.path.join(directory,Sample,Sample + '_particle_images',particle_name),particle)

for x in np.unique(subset_115x.file_name):
	photo = io.imread(os.path.join(image_directory,Sample,Sample+'_115x_images',x), plugin= 'pil')
	photo_subset_115x = subset_115x.loc[subset_115x.file_name == x]
	for p in photo_subset_115x.index:
		particle = photo[int(photo_subset_115x.loc[p].Top):int(photo_subset_115x.loc[p].Bottom),int(photo_subset_115x.loc[p].Left):int(photo_subset_115x.loc[p].Right)]
		particle_name1 = x.split('.jpg')
		particle_name = particle_name1[0] + '_' + str(int(photo_subset_115x.loc[p].Number)) + '.jpg'
		io.imsave(os.path.join(directory,Sample,Sample + '_particle_images',particle_name),particle)
		
		
Measured_particles_allMagnifications = pd.concat([subset_7x,subset_20x,subset_40x,subset_115x])
Measured_particles_allMagnifications.index = np.arange(0,len(Measured_particles_allMagnifications.Number))

particle_image_names = []

for x in Measured_particles_allMagnifications.index:
	filename = Measured_particles_allMagnifications.loc[x].file_name
	filename1=filename.split('.jpg')
	particle_image_name = filename1[0] + '_' + str(int(Measured_particles_allMagnifications.loc[x].Number)) + '.jpg'
	particle_image_names.append(particle_image_name)

Measured_particles_allMagnifications['particle_image_name'] = particle_image_names

Measured_particles_allMagnifications.to_csv(os.path.join(directory,Sample,'Measured_particles_all-magnifications.csv'))
```
