## Supplemental File 1 for "A visual tour of carbon export by sinking particles": ID_from_folders.py

```
#!/Users/colleen/anaconda3/bin/python3

#This code looks in a directory containing subfolders that are named by particle types.
#The particle image filenames within each subfolder are found in the particle csv file and assigned the subfolder name (particle ID) in the csv.
#A new csv file is saved that containes the particle IDs.

import pandas as pd
import numpy as np
import math, os, sys

##Manually currated ID data from images in directories

path = sys.argv[1]
csv_file = sys.argv[2]

data = pd.read_csv(csv_file)

data['ID'] = np.zeros(len(data['Number']))   

categories = os.listdir(path)
for cat in categories:
	if os.path.isdir(os.path.join(path,cat)):
		image_list = os.listdir(os.path.join(path,cat))
		for image in image_list:
			data['ID'].loc[data['particle_image_name']==image] = cat

newname = csv_file.split('.csv')
data.to_csv(newname[0] +'_v2.csv')
```
