## Supplemental File 1 for "A visual tour of carbon export by sinking particles": Calculate_fluxes_Endeavor.py

```
#!/Users/colleen/anaconda3/bin/python3

#This code calculates the number flux of each sinking particle category.  
#Input csv file is generated from previous steps.  First, image processing scripts generate a csv file of all particles detected.
#Second, csv files of detected particles from multiple magnifications are concatenated.
#Third, identities are assigned to each particle in the csv file.
#This concatenated csv file containing particle identities is the input into this script that calculates fluxes.


from scipy import optimize
import pandas as pd
import numpy as np
import math, os, sys
import matplotlib.pyplot as plt


##Calculate fluxes, print graphs, save a csv file with flux data including all particle types

input_csv = sys.argv[1]
sample_name = sys.argv[2]
time = float(sys.argv[3])
output_directory = sys.argv[4]

data = pd.read_csv(input_csv)

##scale of pixels per micron: 7x = 0.1268 , 20x = 0.364 , 50x = 0.928 , 115x = 2.179
##scale is in meters squared##
EN_img_area_7x = (1388/65000) * (1038/65000)
EN_img_area_20x = (1388/187000) * (1038/187000)
EN_img_area_40x = (1388/375000) * (1038/375000)
EN_img_area_80x = (1388/708000) * (1038/708000)
EN_img_area_115x = (1388/1116000) * (1038/1116000)

#define the size bins##
bins = []
for x in np.arange(0,13.5,0.5):
    bin = 2**x
    bins.append(bin)

bin_mids = []
for y in np.arange(0,len(bins)-1):
    mid = bins[y] + (bins[y+1]-bins[y])/2
    bin_mids.append(mid)

bin_width = []
for z in np.arange(0,len(bins)-1):
    width = (bins[z+1]-bins[z])
    bin_width.append(width)

##Add magnification column to the dataframe

def add_magnification(data):
    magnification_list = []
    for x in data.file_name:
        name = x.split('_')
        magnification_list.append(name[2])
    
    data['magnification'] = magnification_list
    return data

data = add_magnification(data)

##Remove noise and define sinking particles
def sinking_particles(data):
    data_good = data[data.ID != 'unidentifiable']
    data_good = data_good[data_good.ID != 'fiber']
    data_good = data_good[data_good.ID != 'copepod']
    data_good = data_good[data_good.ID != 'amphipod']
    data_good = data_good[data_good.ID != 'pteropod']
    data_good = data_good[data_good.ID != 'zooplankton_part']
    data_good = data_good[data_good.ID != 'zooplankton']
    return data_good


data_sinking = sinking_particles(data)

#Calculate particle flux at each magnification and display to identify best bin sizes for each magnification
def Particle_flux(data,time, particle_type=None,img_area_7x=EN_img_area_7x, img_area_20x=EN_img_area_20x,img_area_40x=EN_img_area_40x,img_area_80x = EN_img_area_80x, img_area_115x=EN_img_area_115x,bins=bins, bin_mids=bin_mids, bin_width=bin_width):
    data_7x = data[data.magnification == '7x']
    data_20x = data[data.magnification == '20x']
    data_40x = data[data.magnification == '40x']
    data_80x = data[data.magnification == '80x']
    data_115x = data[data.magnification == '115x']

    fov_7x = int(np.max(data_7x.fov))
    fov_20x = int(np.max(data_20x.fov))
    fov_40x = int(np.max(data_40x.fov))
    try:
        fov_80x = int(np.max(data_80x.fov))
    except:
        fov_115x = int(np.max(data_115x.fov))

    if particle_type != None:
        data = data.loc[data.ID == particle_type]
        data_7x = data[data.magnification == '7x']
        data_20x = data[data.magnification == '20x']
        data_40x = data[data.magnification == '40x']
        try:
            if isinstance(fov_80x, int):
                data_80x = data[data.magnification == '80x']
        except:
            if isinstance(fov_115x, int):
                data_115x = data[data.magnification == '115x']

    histo_7x = np.histogram(data_7x.ESD,bins=bins)
    flux_7x = histo_7x[0]/fov_7x/img_area_7x/time  #in unit of particles per square meter per day
    flux_7x_error = np.sqrt(histo_7x[0])/fov_7x/img_area_7x/time
    number_counted_7x = histo_7x[0]
    result = pd.DataFrame(np.stack((bin_mids,bin_width,flux_7x,flux_7x_error,number_counted_7x),-1),columns=('bin_mids','bin_width','flux_7x','flux_error_7x','number_counted_7x'))

    histo_20x = np.histogram(data_20x.ESD,bins=bins)
    flux_20x = histo_20x[0]/fov_20x/img_area_20x/time  #in unit of particles per square meter per day
    flux_20x_error = np.sqrt(histo_20x[0])/fov_20x/img_area_20x/time
    number_counted_20x = histo_20x[0]
    result_20x = pd.DataFrame(np.stack((flux_20x,flux_20x_error,number_counted_20x),-1),columns=('flux_20x','flux_error_20x','number_counted_20x'))
    
    histo_40x = np.histogram(data_40x.ESD,bins=bins)
    flux_40x = histo_40x[0]/fov_40x/img_area_40x/time  #in unit of particles per square meter per day
    flux_40x_error = np.sqrt(histo_40x[0])/fov_40x/img_area_40x/time
    number_counted_40x = histo_40x[0]
    result_40x = pd.DataFrame(np.stack((flux_40x,flux_40x_error,number_counted_40x),-1),columns=('flux_40x','flux_error_40x','number_counted_40x'))

    try:
        histo_80x = np.histogram(data_80x.ESD,bins=bins)
        flux_80x = histo_80x[0]/fov_80x/img_area_80x/time  #in unit of particles per square meter per day
        flux_80x_error = np.sqrt(histo_80x[0])/fov_80x/img_area_80x/time
        number_counted_80x = histo_80x[0]
        result_80x = pd.DataFrame(np.stack((flux_80x,flux_80x_error,number_counted_80x),-1),columns=('flux_80x','flux_error_80x','number_counted_80x'))

    except:
        histo_115x = np.histogram(data_115x.ESD,bins=bins)
        flux_115x = histo_115x[0]/fov_115x/img_area_115x/time  #in unit of particles per square meter per day
        flux_115x_error = np.sqrt(histo_115x[0])/fov_115x/img_area_115x/time
        number_counted_115x = histo_115x[0]
        result_115x = pd.DataFrame(np.stack((flux_115x,flux_115x_error,number_counted_115x),-1),columns=('flux_115x','flux_error_115x','number_counted_115x'))

    result['flux_20x'] =  flux_20x
    result['flux_error_20x'] =  flux_20x_error
    result['number_counted_20x'] =  number_counted_20x
    result['flux_40x'] =  flux_40x
    result['flux_error_40x'] =  flux_40x_error
    result['number_counted_40x'] =  number_counted_40x
    try:
        result['flux_80x'] =  flux_80x
        result['flux_error_80x'] =  flux_80x_error
        result['number_counted_80x'] =  number_counted_80x
    except:
        result['flux_115x'] =  flux_115x
        result['flux_error_115x'] =  flux_115x_error
        result['number_counted_115x'] =  number_counted_115x

    return result     
#Figure to show how flux was detected at each magnification (sanity checking)
data_flux_allMags = Particle_flux(data_sinking,time)

plt.figure(figsize=([5,8]))
plt.axis([10,10000,0.01,1000000])
plt.loglog(data_flux_allMags.bin_mids[data_flux_allMags.flux_7x!=0],data_flux_allMags.flux_7x[data_flux_allMags.flux_7x!=0]/data_flux_allMags.bin_width[data_flux_allMags.flux_7x!=0],marker = 'o',ms=3,lw=2, label= '7x')
plt.loglog(data_flux_allMags.bin_mids[data_flux_allMags.flux_20x!=0],data_flux_allMags.flux_20x[data_flux_allMags.flux_20x!=0]/data_flux_allMags.bin_width[data_flux_allMags.flux_20x!=0], lw=2, marker = 'o',ms=3,color = 'green', label = '20x')
plt.loglog(data_flux_allMags.bin_mids[data_flux_allMags.flux_40x!=0],data_flux_allMags.flux_40x[data_flux_allMags.flux_40x!=0]/data_flux_allMags.bin_width[data_flux_allMags.flux_40x!=0], lw=2,marker = 'o',ms=3,color = 'orange', label ='40x')
try:
	plt.loglog(data_flux_allMags.bin_mids[data_flux_allMags.flux_115x!=0],data_flux_allMags.flux_115x[data_flux_allMags.flux_115x!=0]/data_flux_allMags.bin_width[data_flux_allMags.flux_115x!=0], lw=2,marker = 'o',ms=3,color = 'red', label ='115x')
except:
	plt.loglog(data_flux_allMags.bin_mids[data_flux_allMags.flux_80x!=0],data_flux_allMags.flux_80x[data_flux_allMags.flux_80x!=0]/data_flux_allMags.bin_width[data_flux_allMags.flux_80x!=0], lw=2,marker = 'o',ms=3,color = 'red', label ='80x')

plt.legend(frameon=False,fontsize='small')
plt.title('Particle flux at all magnifications')
plt.savefig(os.path.join(output_directory,sample_name+'_Flux_all_magnifications.pdf'))

#Use flux data from only the most optimal magnification for a given size bin.
def flux_combined(data,threshold_7x, threshold_20x, threshold_40x, threshold_80_or_115x):
    data.bin_mids=data.bin_mids.astype('int')
    subset_7x = data.loc[data.bin_mids >= threshold_7x]
    subset_20xa = data.loc[data.bin_mids >= threshold_20x]
    subset_20x = subset_20xa.loc[subset_20xa.bin_mids < threshold_7x]
    subset_40xa = data.loc[data.bin_mids >= threshold_40x]
    subset_40x = subset_40xa.loc[subset_40xa.bin_mids < threshold_20x]
    subset_80_or_115xa = data.loc[data.bin_mids >= threshold_80_or_115x]
    subset_80_or_115x = subset_80_or_115xa.loc[subset_80_or_115xa.bin_mids < threshold_40x]

    PSD_flux_all= pd.DataFrame()
    try:
        flux_all = np.concatenate((subset_80_or_115x.flux_80x,subset_40x.flux_40x,subset_20x.flux_20x,subset_7x.flux_7x))
    except:
        flux_all = np.concatenate((subset_80_or_115x.flux_115x,subset_40x.flux_40x,subset_20x.flux_20x,subset_7x.flux_7x))
    try:
        counts_all = np.concatenate((subset_80_or_115x.number_counted_80x,subset_40x.number_counted_40x,subset_20x.number_counted_20x,subset_7x.number_counted_7x))
    except:
        counts_all = np.concatenate((subset_80_or_115x.number_counted_115x,subset_40x.number_counted_40x,subset_20x.number_counted_20x,subset_7x.number_counted_7x))
    try:
        flux_error_all = np.concatenate((subset_80_or_115x.flux_error_80x,subset_40x.flux_error_40x,subset_20x.flux_error_20x,subset_7x.flux_error_7x))
    except:
        flux_error_all = np.concatenate((subset_80_or_115x.flux_error_115x,subset_40x.flux_error_40x,subset_20x.flux_error_20x,subset_7x.flux_error_7x))
    bin_mids_all = np.concatenate((subset_80_or_115x.bin_mids,subset_40x.bin_mids,subset_20x.bin_mids,subset_7x.bin_mids))
    bin_width_all = np.concatenate((subset_80_or_115x.bin_width,subset_40x.bin_width,subset_20x.bin_width,subset_7x.bin_width))
    PSD_flux_all['bin_mids'] = bin_mids_all
    PSD_flux_all['bin_width'] = bin_width_all
    PSD_flux_all['flux'] = flux_all
    PSD_flux_all['flux_error'] = flux_error_all
    PSD_flux_all['number_counted'] = counts_all

    return PSD_flux_all

#Create new dataframe of total flux including only optimal magnifications
data_flux_combined = flux_combined(data_flux_allMags,437,154,109,11)

#Calculate the PSD slope using data >54 um and 54 um size data as the reference value
def PSD(var, bin_mid, normalized_flux,norm_bin_value):
    norm_flux_data = normalized_flux.loc[normalized_flux>0]
    bin_mid_data = bin_mid.loc[normalized_flux>0]
    normalization_scalar = norm_flux_data.loc[bin_mid_data == norm_bin_value]
    norm_flux_data = norm_flux_data.loc[bin_mid_data >= norm_bin_value]
    N = normalization_scalar.values * (bin_mid_data/norm_bin_value)**var
    Z = np.sum((np.log(N)-np.log(norm_flux_data))**2)
    return Z
    
data_PSDslope = optimize.minimize(PSD, [-3], args=(data_flux_combined.bin_mids, data_flux_combined.flux/data_flux_combined.bin_width,54))

#Calculate the flux of each particle type
data_agg_flux_allMags = Particle_flux(data_sinking,time,'aggregate')
data_agg_flux_combined = flux_combined(data_agg_flux_allMags,437,154,109,11)
data_long_flux_allMags = Particle_flux(data_sinking,time,'long_fecal_pellet')
data_long_flux_combined = flux_combined(data_long_flux_allMags,437,154,109,11)
data_large_flux_allMags = Particle_flux(data_sinking,time,'large_loose_pellet')
data_large_flux_combined = flux_combined(data_large_flux_allMags,437,154,109,11)
data_short_flux_allMags = Particle_flux(data_sinking,time,'short_pellet')
data_short_flux_combined = flux_combined(data_short_flux_allMags,437,154,109,11)
data_mini_flux_allMags = Particle_flux(data_sinking,time,'mini_pellet')
data_mini_flux_combined = flux_combined(data_mini_flux_allMags,437,154,109,11)
data_dense_flux_allMags = Particle_flux(data_sinking,time,'dense_detritus')
data_dense_flux_combined = flux_combined(data_dense_flux_allMags,437,154,109,11)
data_phyto_flux_allMags = Particle_flux(data_sinking,time,'phytoplankton')
data_phyto_flux_combined = flux_combined(data_phyto_flux_allMags,437,154,109,11)
data_foraminifera_flux_allMags = Particle_flux(data_sinking,time,'foraminifera')
data_foraminifera_flux_combined = flux_combined(data_foraminifera_flux_allMags,437,154,109,11)
data_rhizaria_flux_allMags = Particle_flux(data_sinking,time,'rhizaria')
data_rhizaria_flux_combined = flux_combined(data_rhizaria_flux_allMags,437,154,109,11)
data_copepod_flux_allMags = Particle_flux(data,time,'copepod')
data_copepod_flux_combined = flux_combined(data_copepod_flux_allMags,437,154,109,11)
data_amphipod_flux_allMags = Particle_flux(data,time,'amphipod')
data_amphipod_flux_combined = flux_combined(data_amphipod_flux_allMags,437,154,109,11)
data_zooplankton_flux_allMags = Particle_flux(data,time,'zooplankton')
data_zooplankton_flux_combined = flux_combined(data_zooplankton_flux_allMags,437,154,109,11)
data_zooplankton_part_flux_allMags = Particle_flux(data,time,'zooplankton_part')
data_zooplankton_part_flux_combined = flux_combined(data_zooplankton_part_flux_allMags,437,154,109,11)
data_fiber_flux_allMags = Particle_flux(data,time,'fiber')
data_fiber_flux_combined = flux_combined(data_fiber_flux_allMags,437,154,109,11)
data_unidentifiable_flux_allMags = Particle_flux(data,time,'unidentifiable')
data_unidentifiable_flux_combined = flux_combined(data_unidentifiable_flux_allMags,437,154,109,11)
data_pteropod_flux_allMags = Particle_flux(data,time,'pteropod')
data_pteropod_flux_combined = flux_combined(data_pteropod_flux_allMags,437,154,109,11)
data_salp_pellet_flux_allMags = Particle_flux(data,time,'salp_pellet')
data_salp_pellet_flux_combined = flux_combined(data_salp_pellet_flux_allMags,437,154,109,11)


data_all_flux_data = data_flux_combined.merge(data_agg_flux_combined,on='bin_mids',how='left')
data_all_flux_data=data_all_flux_data.rename(index=str, columns={"flux_x": "flux", "bin_width_x":"bin_width", "flux_error_x": "flux_uncertainty","number_counted_x":"number_counted","flux_y":"aggregate_flux","flux_error_y":"aggregate_flux_uncertainty","number_counted_y":"aggregate_number_counted"})
data_all_flux_data=data_all_flux_data.merge(data_long_flux_combined,on='bin_mids',how='left')
data_all_flux_data=data_all_flux_data.rename(index=str, columns={"flux_x": "flux", "bin_width_x":"bin_width","number_counted_x":"number_counted","flux_y":"long_fp_flux","flux_error":"long_fp_flux_uncertainty","number_counted_y":"long_fp_number_counted"})
data_all_flux_data=data_all_flux_data.merge(data_dense_flux_combined,on='bin_mids',how='left')
data_all_flux_data=data_all_flux_data.rename(index=str, columns={"flux_x": "flux", "bin_width_x":"bin_width","number_counted_x":"number_counted","flux_y":"dense_detritus_flux","flux_error":"dense_detritus_flux_uncertainty","number_counted_y":"dense_detritus_number_counted"})
data_all_flux_data=data_all_flux_data.merge(data_large_flux_combined,on='bin_mids',how='left')
data_all_flux_data=data_all_flux_data.rename(index=str, columns={"flux_x": "flux", "bin_width_x":"bin_width","number_counted_x":"number_counted","flux_y":"large_loose_flux","flux_error":"large_loose_flux_uncertainty","number_counted_y":"large_loose_number_counted"})
data_all_flux_data=data_all_flux_data.merge(data_short_flux_combined,on='bin_mids',how='left')
data_all_flux_data=data_all_flux_data.rename(index=str, columns={"flux_x": "flux", "bin_width_x":"bin_width","number_counted_x":"number_counted","flux_y":"short_fp_flux","flux_error":"short_fp_flux_uncertainty","number_counted_y":"short_fp_number_counted"})
data_all_flux_data=data_all_flux_data.merge(data_mini_flux_combined,on='bin_mids',how='left')
data_all_flux_data=data_all_flux_data.rename(index=str, columns={"flux_x": "flux", "bin_width_x":"bin_width","number_counted_x":"number_counted","flux_y":"mini_pellet_flux","flux_error":"mini_pellet_flux_uncertainty","number_counted_y":"mini_pellet_number_counted"})
data_all_flux_data=data_all_flux_data.merge(data_salp_pellet_flux_combined,on='bin_mids',how='left')
data_all_flux_data=data_all_flux_data.rename(index=str, columns={"flux_x": "flux", "bin_width_x":"bin_width","number_counted_x":"number_counted","flux_y":"salp_pellet_flux","flux_error":"salp_pellet_flux_uncertainty","number_counted_y":"salp_pellet_number_counted"})
data_all_flux_data=data_all_flux_data.merge(data_phyto_flux_combined,on='bin_mids',how='left')
data_all_flux_data=data_all_flux_data.rename(index=str, columns={"flux_x": "flux", "bin_width_x":"bin_width","number_counted_x":"number_counted","flux_y":"phyto_flux","flux_error":"phyto_flux_uncertainty","number_counted_y":"phyto_number_counted"})
data_all_flux_data=data_all_flux_data.merge(data_foraminifera_flux_combined,on='bin_mids',how='left')
data_all_flux_data=data_all_flux_data.rename(index=str, columns={"flux_x": "flux", "bin_width_x":"bin_width","number_counted_x":"number_counted","flux_y":"foraminifera_flux","flux_error":"foraminifera_flux_uncertainty","number_counted_y":"foraminifera_number_counted"})
data_all_flux_data=data_all_flux_data.merge(data_rhizaria_flux_combined,on='bin_mids',how='left')
data_all_flux_data=data_all_flux_data.rename(index=str, columns={"flux_x": "flux", "bin_width_x":"bin_width","number_counted_x":"number_counted","flux_y":"rhizaria_flux","flux_error":"rhizaria_flux_uncertainty","number_counted_y":"rhizaria_number_counted"})
data_all_flux_data=data_all_flux_data.merge(data_fiber_flux_combined,on='bin_mids',how='left')
data_all_flux_data=data_all_flux_data.rename(index=str, columns={"flux_x": "flux", "bin_width_x":"bin_width","number_counted_x":"number_counted","flux_y":"fiber_flux","flux_error":"fiber_flux_uncertainty","number_counted_y":"fiber_number_counted"})
data_all_flux_data=data_all_flux_data.merge(data_copepod_flux_combined,on='bin_mids',how='left')
data_all_flux_data=data_all_flux_data.rename(index=str, columns={"flux_x": "flux", "bin_width_x":"bin_width","number_counted_x":"number_counted","flux_y":"copepod_flux","flux_error":"copepod_flux_uncertainty","number_counted_y":"copepod_number_counted"})
data_all_flux_data=data_all_flux_data.merge(data_pteropod_flux_combined,on='bin_mids',how='left')
data_all_flux_data=data_all_flux_data.rename(index=str, columns={"flux_x": "flux", "bin_width_x":"bin_width","number_counted_x":"number_counted","flux_y":"pteropod_flux","flux_error":"pteropod_flux_uncertainty","number_counted_y":"pteropod_number_counted"})
data_all_flux_data=data_all_flux_data.merge(data_amphipod_flux_combined,on='bin_mids',how='left')
data_all_flux_data=data_all_flux_data.rename(index=str, columns={"flux_x": "flux", "bin_width_x":"bin_width","number_counted_x":"number_counted","flux_y":"amphipod_flux","flux_error":"amphipod_flux_uncertainty","number_counted_y":"amphipod_number_counted"})
data_all_flux_data=data_all_flux_data.merge(data_zooplankton_flux_combined,on='bin_mids',how='left')
data_all_flux_data=data_all_flux_data.rename(index=str, columns={"flux_x": "flux", "bin_width_x":"bin_width","number_counted_x":"number_counted","flux_y":"other_zooplankton_flux","flux_error":"other_zooplankton_flux_uncertainty","number_counted_y":"other_zooplankton_number_counted"})
data_all_flux_data=data_all_flux_data.merge(data_zooplankton_part_flux_combined,on='bin_mids',how='left')
data_all_flux_data=data_all_flux_data.rename(index=str, columns={"flux_x": "flux", "bin_width_x":"bin_width","number_counted_x":"number_counted","flux_y":"zooplankton_part_flux","flux_error":"zooplankton_part_flux_uncertainty","number_counted_y":"zooplankton_part_number_counted"})
data_all_flux_data=data_all_flux_data.merge(data_unidentifiable_flux_combined,on='bin_mids',how='left')
data_all_flux_data=data_all_flux_data.rename(index=str, columns={"flux_x": "flux", "bin_width_x":"bin_width","number_counted_x":"number_counted","flux_y":"unidentifiable_flux","flux_error":"unidentifiable_flux_uncertainty","number_counted_y":"unidentifiable_number_counted"})

data_all_flux_data=data_all_flux_data.drop('bin_width_y',axis=1)
data_all_flux_data.to_csv(os.path.join(output_directory,sample_name+"_all_flux_data.csv"))

data_mids = data_flux_combined.bin_mids
data_mids.iloc[0]=13.4
data_agg_top = data_agg_flux_combined.flux/data_flux_combined.flux
data_agg_top=data_agg_top.fillna(0)
data_dense_top = data_dense_flux_combined.flux/data_flux_combined.flux
data_dense_top = data_dense_top.fillna(0)
data_long_top = data_long_flux_combined.flux/data_flux_combined.flux
data_long_top=data_long_top.fillna(0)
data_large_top = data_large_flux_combined.flux/data_flux_combined.flux
data_large_top=data_large_top.fillna(0)
data_salp_top = data_salp_pellet_flux_combined.flux/data_flux_combined.flux
data_salp_top=data_salp_top.fillna(0)
data_short_top = data_short_flux_combined.flux/data_flux_combined.flux
data_short_top = data_short_top.fillna(0)
data_mini_top = data_mini_flux_combined.flux/data_flux_combined.flux
data_mini_top = data_mini_top.fillna(0)
data_foraminifera_top = data_foraminifera_flux_combined.flux/data_flux_combined.flux
data_foraminifera_top = data_foraminifera_top.fillna(0)
data_phyto_top = data_phyto_flux_combined.flux/data_flux_combined.flux
data_phyto_top = data_phyto_top.fillna(0)
data_rhizaria_top = data_rhizaria_flux_combined.flux/data_flux_combined.flux
data_rhizaria_top = data_rhizaria_top.fillna(0)

fig = plt.figure(figsize=(6,4))
ax1 = fig.add_subplot(111)
ax1.semilogx()
ax1.set_xlabel('ESD (um)')
ax1.set_ylabel('fraction of number flux)')
ax1.set_title(sample_name)
ax1.set_xlim([10,100000])
ax1.bar(data_mids,data_agg_top, width=data_flux_combined.bin_width, color = 'navy',label='aggregate')
ax1.bar(data_mids,data_dense_top,bottom=data_agg_top, width=data_flux_combined.bin_width, color = 'gold',label='dense detritus')
ax1.bar(data_mids,data_salp_top,bottom=data_agg_top+data_dense_top, width=data_flux_combined.bin_width, color = 'orange',label='salp fp')
ax1.bar(data_mids,data_large_top,bottom=data_salp_top+data_agg_top+data_dense_top, width=data_flux_combined.bin_width, color = 'crimson',label='large loose fp')
ax1.bar(data_mids,data_long_top,bottom=data_salp_top+data_large_top+data_agg_top+data_dense_top, width=data_flux_combined.bin_width, color = 'red',label='long fp')
ax1.bar(data_mids,data_short_top,bottom=data_salp_top+data_large_top+data_long_top+data_agg_top+data_dense_top, width=data_flux_combined.bin_width, color = 'grey',label='short fp')
ax1.bar(data_mids,data_mini_top,bottom=data_salp_top+data_large_top+data_short_top+data_long_top+data_agg_top+data_dense_top, width=data_flux_combined.bin_width, color = 'deepskyblue',label='mini pellet')
ax1.bar(data_mids,data_foraminifera_top,bottom=data_salp_top+data_large_top+data_mini_top+data_short_top+data_long_top+data_agg_top+data_dense_top, width=data_flux_combined.bin_width, color = 'black',label='foraminifera')
ax1.bar(data_mids,data_phyto_top,bottom=data_salp_top+data_large_top+data_foraminifera_top+data_mini_top+data_short_top+data_long_top+data_agg_top+data_dense_top, width=data_flux_combined.bin_width, color = 'green',label='phytoplankton')
ax1.bar(data_mids,data_rhizaria_top,bottom=data_salp_top+data_large_top+data_phyto_top+data_foraminifera_top+data_mini_top+data_short_top+data_long_top+data_agg_top+data_dense_top, width=data_flux_combined.bin_width, color = 'lawngreen',label='rhizaria')
ax1.text(3000,0.2,'slope =' + str(data_PSDslope.x[0])[0:6],color = 'black',fontsize=10)
plt.legend(loc=1,frameon=False,fontsize='x-small')
ax2 = ax1.twinx()
ax2.loglog(data_mids[data_flux_combined.flux!=0],data_flux_combined.flux[data_flux_combined.flux!=0]/data_flux_combined.bin_width[data_flux_combined.flux!=0],lw=2,color='white')
ax2.set_ylabel('number flux (m-2 d-1 um-1)')
plt.tight_layout()
plt.savefig(os.path.join(output_directory,sample_name + ' PSD of particle types.pdf'))
```
